# STK19 positions TFIIH for cell-free transcription-coupled DNA repair

**DOI:** 10.1101/2024.07.22.604623

**Authors:** Tycho E.T. Mevissen, Maximilian Kümmecke, Ernst W. Schmid, Lucas Farnung, Johannes C. Walter

**Affiliations:** Department of Biological Chemistry and Molecular Pharmacology, Blavatnik Institute, Harvard Medical School, Boston, MA 02115, USA; Department of Cell Biology, Blavatnik Institute, Harvard Medical School, Boston, MA 02115, USA; Howard Hughes Medical Institute

## Abstract

In transcription-coupled repair, stalled RNA polymerase II (Pol II) is recognized by CSB and CRL4^CSA^, which co-operate with UVSSSA and ELOF1 to recruit TFIIH for nucleotide excision repair (TC-NER). To explore the mechanism of TC-NER, we recapitulated this reaction *in vitro*. When a plasmid containing a site-specific lesion is transcribed in frog egg extract, error-free repair is observed that depends on CSB, CRL4^CSA^, UVSSA, and ELOF1. Repair also depends on STK19, a factor previously implicated in transcription recovery after UV exposure. A 1.9 Å cryo-electron microscopy structure shows that STK19 joins the TC-NER complex by binding CSA and the RPB1 subunit of Pol II. Furthermore, AlphaFold predicts that STK19 interacts with the XPD subunit of TFIIH, and disrupting this interface impairs cell-free repair. Molecular modeling suggests that STK19 positions TFIIH ahead of Pol II for lesion verification. In summary, our analysis of cell-free TC-NER suggests that STK19 couples RNA polymerase II stalling to downstream repair events.

## INTRODUCTION

Our cells contain numerous mechanisms to repair DNA damage that is continually generated by diverse endogenous and exogenous agents. A particularly versatile pathway is nucleotide excision repair (NER), which removes bulky DNA adducts regardless of their chemical structure.^1,2^ In global genome (GG)-NER, which can in principle operate at any locus, Xeroderma pigmentosum group protein C (XPC) in complex with RAD23B and Centrin 2 (CETN2) recognizes the distortion in DNA structure created by bulky lesions. This heterotrimeric complex then recruits TFIIH, whose XPB ATPase subunit unwinds DNA surrounding the lesion, and whose XPD ATPase subunit searches one strand for the presence of DNA damage.^3^ If a lesion is detected, TFIIH recruits the downstream repair machinery, including two structure-specific endonucleases, XPF-ERCC1 and XPG, which incise the damaged strand on either side of the lesion. The damaged oligonucleotide dissociates from DNA, and gap filling completes the repair reaction. GG-NER has been reconstituted with purified components and is therefore relatively well-understood.^2,4,5^

Almost 40 years ago, the Hanawalt group discovered that DNA damage located in the transcribed strand of a gene is preferentially repaired by NER, leading to the concept of transcription-coupled (TC)-NER.^6–8^ In this mechanism, DNA damage is sensed by RNA polymerase II (Pol II) stalling instead of by XPC-RAD23B-CETN2. Four factors have been identified that couple Pol II stalling to TFIIH recruitment and the same downstream repair steps that operate in GG-NER.^9^ The first is CSB, which is mutated in a human neurodegenerative progeroid syndrome called Cockayne syndrome. CSB is a SWI/SNF-type ATPase that binds on the upstream side of stalled Pol II and attempts to push it past obstacles.^10,11^ If the obstacle is insurmountable, as seen for many DNA lesions, CSB recruits the CRL4^CSA^ E3 ubiquitin ligase whose substrate receptor CSA links to a CUL4 scaffold via DDB1. CSB recruits CRL4^CSA^ via a short CSA-interacting motif (CIM) that binds directly to CSA.^12^ CRL4^CSA^ attaches ubiquitin to lysine 1268 of RPB1, the largest subunit of Pol II, and this modification is required for TFIIH recruitment via an unknown mechanism. The third TC-NER factor that is also a transcription elongation factor, ELOF1, interacts with Pol II and CRL4^CSA^, and it is required for efficient Pol II polyubiquitination.^13–15^ Finally, UV sensitivity syndrome protein A (UVSSA), is recruited to stalled Pol II via a direct interaction with CSA,^11^ and UVSSA binding and Pol II ubiquitination appear to be interdependent.^16^ In turn, UVSSA interacts with and is essential to recruit TFIIH to the repair complex via direct binding to the p62 subunit.^12,17^ However, TFIIH interacts with an unstructured region of UVSSA (the TFIIH-interacting region, TIR), leaving open the question of how TFIIH is properly positioned ahead of Pol II to recognize the damaged strand. Thus, the mechanism by which Pol II stalling is coupled to downstream repair events remains incompletely understood.

Serine threonine kinase 19 (STK19) was nominated by several groups as a possible TC-NER factor. Despite its name, STK19 bears no resemblance to protein kinases, and the purified protein has no detectable kinase activity.^18,19^ A chemogenomic CRISPR screen revealed that STK19-deficient cells are sensitive to the alkylating agent illudin S, as seen for other TC-NER factors.^20^ Moreover, Boeing *et al*. showed that STK19 is required for the recovery of transcription after cellular exposure to UV light.^21^ These observations are consistent with a role for STK19 in TC-NER but might also indicate a specific function in transcription restart. Thus, whether STK19 is a core TC-NER factor and what role it plays in the response to DNA damage are unanswered questions. A full understanding of TC-NER requires biochemical and structural analysis. To recapitulate TC-NER, Egly and colleagues stalled Pol II at a cisplatin lesion and then mixed it with all the GG-NER factors except XPC.^22^ They found that the addition of CSB to this system promoted a low level (∼1%) of lesion excision. However, unlike in cells,^12^ the recruitment of TFIIH to the lesion was CSB-independent, and the reaction presumably did not contain CRL4^CSA^, UVSSA, or ELOF1, suggesting it reflected a partial TC-NER process. More recently, impressive progress in the structural elucidation of TC-NER complexes has been made. Thus, Pol II complexes containing CSB, CRL4^CSA^, UVSSA, and ELOF1 have been determined by cryo-EM.^11,15,23^ However, the transition to downstream repair events has not been structurally resolved. Thus, a full mechanistic understanding of TC-NER is still lacking.

Given that frog egg extracts recapitulate numerous DNA repair pathways including GG-NER,^24^ we asked whether they might also support TC-NER. To this end, we first recapitulated efficient and inducible *in vitro* transcription in *X. laevis* egg extracts using a plasmid with a strong promoter. Placement of a cisplatin intrastrand crosslink in the template strand downstream of the promoter led to Pol II stalling but no TC-NER. When we supplemented the extract with recombinant CSB, CRL4^CSA^, UVSSA, ELOF1, and STK19, we observed lesion repair that was independent of XPC and abolished by the Pol II inhibitor α-amanitin. Repair depended on all five of the above factors, establishing egg extracts as a system that supports *bona fide* TC-NER *in vitro* and indicating that STK19 is an essential TC-NER factor. To understand how STK19 promotes repair, we used AlphaFold-Multimer and single-particle cryo-EM to elucidate its interaction with the TC-NER machinery. Together with structure-function analyses, our study shows that STK19 is an integral component of the TC-NER complex that interacts with CRL4^CSA^ and RPB1. Molecular modeling and site-directed mutagenesis further suggests that STK19 positions TFIIH in front of the TC-NER complex such that the damaged strand is positioned for lesion verification by the XPD helicase. Our work suggests that STK19 forms the linchpin between lesion-stalled Pol II and downstream repair events.

## RESULTS

### Inducible transcription in frog egg extracts

To recapitulate cell-free TC-NER, we first sought to achieve efficient and inducible transcription in frog egg extracts. To this end, we constructed a plasmid containin a strong basal promoter flanked by GAL4 upstream activating sequences (UAS; Figure 1A). The plasmid was added to a concentrated nucleoplasmic extract (NPE) derived from frog eggs that was also supplemented with TBP, the transcriptional activator GAL4-VP64, and radioactive UTP (Figure 1A). Unlike a total egg lysate, NPE supported transcription (Figure S1A, lanes 1-3 and 7-9) that was greatly stimulated by GAL4-VP64 and TBP (Figure S1A, lanes 10-12). Transcription efficiency was further enhanced via the use of a synthetic super core promoter (Figure S1B) and molecular crowding agents (Figure S1C). When we combined all the above features, transcription greatly exceeded the level observed from an endogenous promoter in NPE^25^ (Figure S1D). Other properties of this inducible, cell-free transcription system will be described elsewhere.

**Figure 1.**
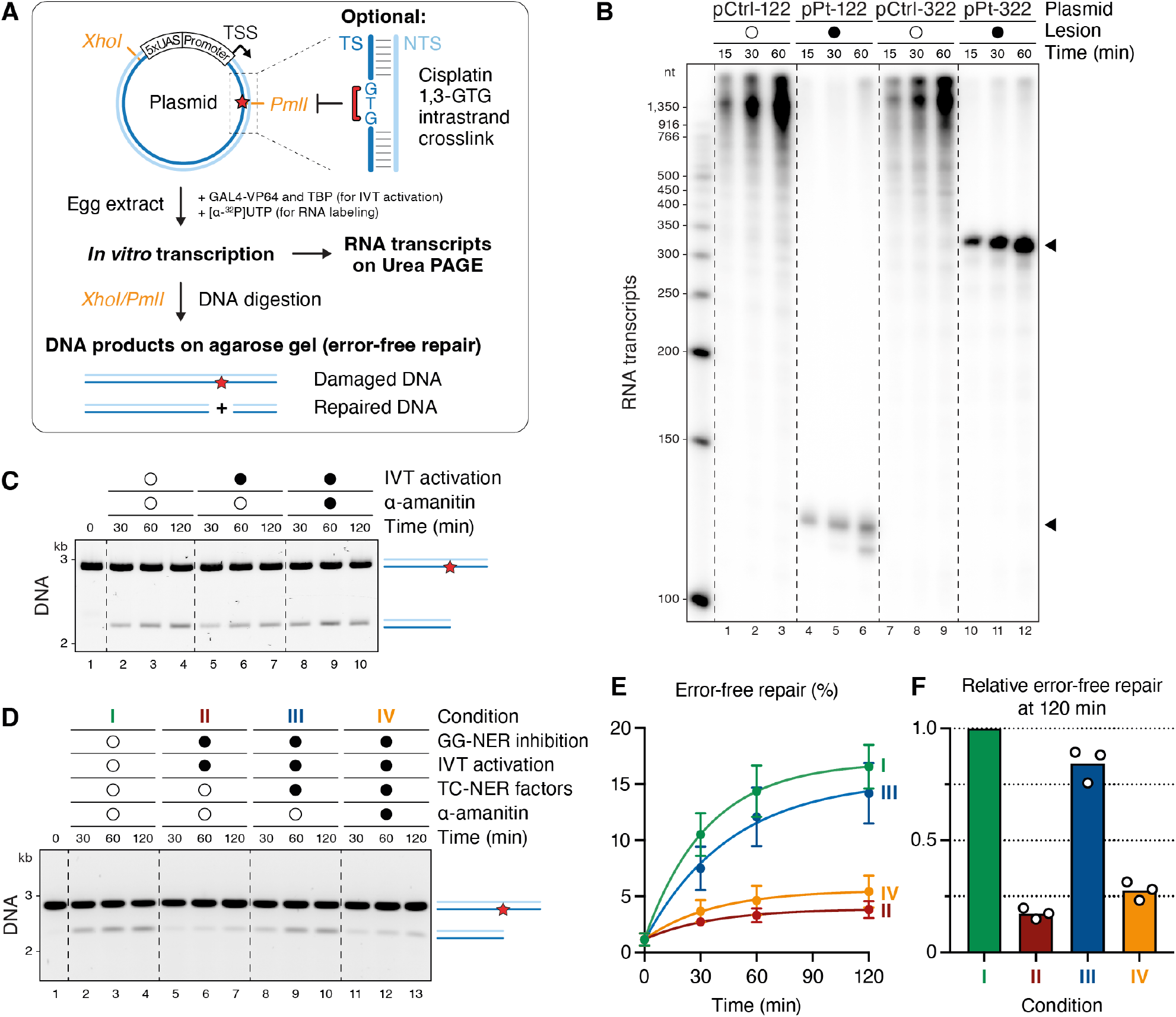
Transcription-coupled DNA repair in egg extracts. **(A)**Generic schematic of the plasmids used for *in vitro* transcription and transcription-coupled DNA repair (including ones containing adenovirus major late and SCP2 promoters). The workflow for analyzing RNA transcripts and error-free DNA repair is outlined. UAS, upstream activation sequence; TSS, transcription start site; TS, template strand; NTS, non-template strand; IVT, *in vitro* transcription. **(B)** Plasmids without (pCtrl) or with cisplatin 1,3-GTG intrastrand crosslinks (pPt) positioned 122 bp or 322 bp downstream of the TSS were added to NPE containing GAL4-VP64, TBP, and [α-^32^P]UTP. At the indicated times, RNA was recovered, separated on a Urea-PAGE gel, and subjected to autoradiography. **(C)** pPt-322 was incubated with NPE that was optionally supplemented with IVT activation mix (GAL4-VP64 and TBP) and α-amanitin (2 µM), as indicated (but not [α-^32^P]UTP). DNA was recovered at indicated times, incubated with XhoI and PmlI, separated on an agarose gel, and visualized with SYBR Gold. Appearance of the 2.2 kb restriction fragment indicates restoration of the PmlI site (the smaller, 0.8 kb fragment is not shown). See Figure S1E for inhibitory effect of α-amanitin. **(D)** Same assay as in (C) but using pPt-122. GG-NER was inhibited via addition of an inhibitory XPC antibody, transcription was induced, and the cocktail of TC-NER factors was added, as indicated. **(E)** Quantification of three experiments like the one shown in (D). Error bars represent the SD from the mean. **(F)** Bar graph quantifying the error-free repair relative to condition I (GG-NER) at the 120 minute time point from (E), our standard approach to present the data throughout the paper.

### Cell-free TC-NER in egg extract

Having achieved efficient cell-free transcription, we sought to recapitulate TC-NER *in vitro*. To this end, we placed a cisplatin 1,3-GTG intrastrand crosslink in the transcribed DNA strand 122 or 322 base pairs downstream of the transcription start site (Figure 1A). These lesions induced a potent block to transcription at the expected location (Figure 1B, lanes 4-6 and 10-12). To measure repair, we asked whether a PmlI restriction site that coincides with the crosslink is regenerated (Figure 1A). As shown in Figure 1C, the PmlI site was regenerated in NPE regardless of whether transcription was induced (lanes 2-7), and repair was unaffected by the transcription inhibitor α-amanitin (lanes 8-10; Figure S1E). When we inhibited or depleted the GG-NER factor XPC, repair was greatly reduced (Figure S1F). Thus, naïve NPE only supported GG-NER, even in the presence of transcription activation.

Based on mass spectrometry analysis, egg extracts contain low or undetectable levels of CSB, CSA, UVSSA, ELOF1, and the candidate TC-NER factor, STK19.^26^ Furthermore, western blotting indicated that the concentrations of CSB and CSA are low in the egg and increase during development (Figure S2A). Based on these observations, we hypothesized that the absence of TC-NER in NPE was due to the absence of one or more TC-NER factors in this extract. To test this idea, we expressed recombinant CSB, CRL4^CSA^, ELOF1, UVSSA, and STK19 (all proteins are from *X. laevis* except CSB, which is human; see Figure S2B and its legend), and combined them to make a “TC-NER cocktail.” Strikingly, in extracts that were undergoing transcription and where XPC-dependent GG-NER was inhibited, the addition of this cocktail stimulated repair (Figure 1D, compare conditions II and III; Figures 1E and 1F for quantification). Moreover, repair was returned to basal levels by the addition of α-amanitin (condition IV). The induction of XPC-independent repair that requires transcription and a cocktail of TC-NER factors strongly suggested that NPE can support cell-free TC-NER.

### Cell-free repair requires all canonical TC-NER factors, as well as STK19

To further test whether our cell-free system recapitulates *bona fide* TC-NER, we omitted each of the five proteins from the cocktail. When CSB was omitted, TC-NER was abolished, but CSB alone supported little repair (Figure 2A, conditions II-IV; Figure S2C). Therefore, CSB is necessary but not sufficient to induce TC-NER in NPE. In the absence of each of the other four factors, repair was modestly reduced (Figure 2A, conditions V-VIII; Figures S2D-S2G). This finding suggested that these four factors are individually not required for cell-free TC-NER, or that the endogenous proteins are sufficient to support repair, despite being undetectable in some cases. To distinguish between these possibilities, we depleted each protein from the extract. When CSA was depleted from NPE and omitted from the cocktail, repair was dramatically reduced, and it was restored by the inclusion of CRL4^CSA^ or CSA-DDB1 in the cocktail (see Methods) (Figure 2B, conditions III and IV; Figure S2D). This result shows that CSA-DDB1 is essential for efficient cell-free TC-NER. Similar results were observed for ELOF1 and UVSSA (Figure 2B, conditions V-VIII; Figures S2E and S2F), demonstrating that cell-free repair in NPE requires all four canonical TC-NER factors (CSB, CRL4^CSA^, UVSSA, and ELOF1). Finally, STK19 was required, strongly arguing that it is a core TC-NER protein (Figures 2B and S2G).

We next addressed whether repair in this system requires previously characterized protein-protein interactions and activities. Repair was inhibited when we disrupted the known interaction between CSA and the CIM of CSB (Figures 2C and S2H), the interactions between ELOF1 and both Pol II and CSA (Figures 2D and S2I), or the interactions between UVSSA and its two binding partners CSA and TFIIH (Figures 2E and S2J).^12,13,15,17^ Moreover, repair was blocked by the general cullin inhibitor MLN4924, consistent with CRL4^CSA^ activity being required for TC-NER (Figures 2F and S2K). Finally, using restriction enzymes whose staggered cutting allows differentiation of the two DNA strands (Figure 2G), we verified that cell-free TC-NER involves unscheduled DNA synthesis (UDS) on the damaged template strand (Figure 2H, condition III), as seen for GG-NER (condition I). Altogether, these results show that egg extracts recapitulate all features expected of TC-NER: involvement of CSB, CRL4^CSA^, ELOF1, and UVSSA; known interactions between these factors; cullin E3 ligase activity; and gap filling on the transcribed strand. Furthermore, the data provide strong evidence that STK19 is a core TC-NER factor that acts upstream of error-free repair.

**Figure 2.**
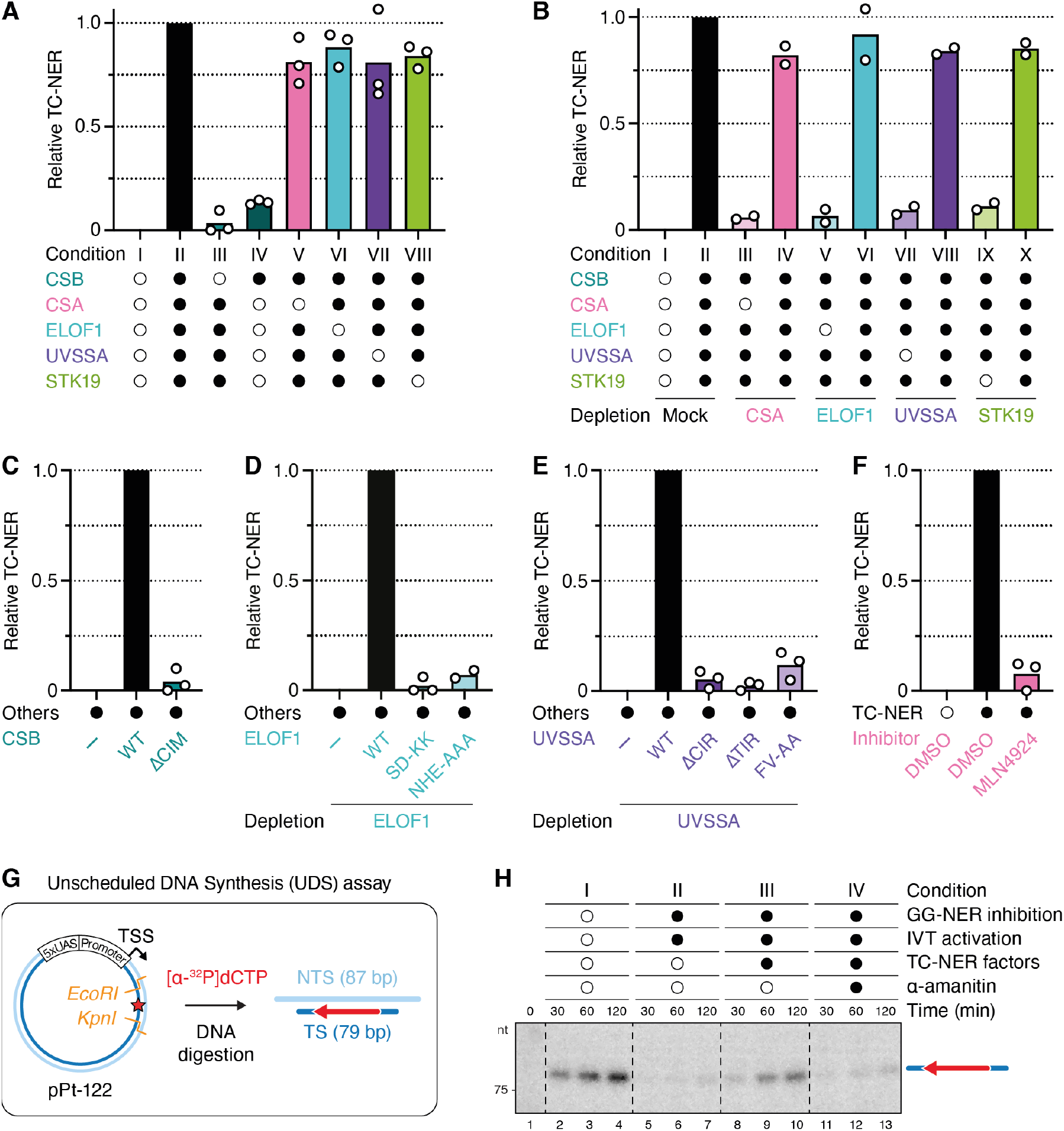
TC-NER in egg extracts requires known proteins, interactions, and STK19. **(A)** Error-free repair assay as described in Figures 1D and 1F, except that GG-NER was inhibited by immunodepletion of XPC from NPE. Open circles indicate factors that were omitted from the TC-NER cocktail. Data of three independent experiments for each condition is plotted. The quantified TC-NER activity for each condition is normalized relative to no repair in condition I and fully active repair in condition II, which are given values of 0 and 1, respectively. Condition II is the standard “TC-NER assay” used in all subsequent experiments. See Figures S2C-S2G for western blots of extracts used. **(B)** TC-NER assay with NPE in which indicated proteins were immunodepleted. Open circles indicate factors omitted from the TC-NER cocktail. See Figures S2D-S2G for western blots of extracts used. **(C)** TC-NER assay comparing recombinant human CSB variants. WT, wild-type; ΔCIM, amino acids 1385-1399 were deleted. See Figure S2H for western blot of extracts used. **(D)** TC-NER assay in ELOF1-depleted NPE. Buffer or the indicated *X. laevis* ELOF1 constructs were added. SD-KK, combination of S72K and D73K to disrupt Pol II binding. NHE-AAA, amino acids N30, H31, and E32 were mutated to alanine to prevent CSA binding. See Figure S2I for western blot of extracts used. **(E)** TC-NER assay in UVSSA-depleted NPE. Buffer or the indicated *X. laevis* UVSSA proteins expressed in wheat germ extract were added back. ΔCIR, amino acids 99-201 were deleted, based on the analogous mutation that disrupts CSA binding in humans.^12^ ΔTIR, residues 416-524 were deleted. FV-AA, residues F425 and V428 were mutated to alanine. See Figure S2J for western blot of extracts used. **(F)** TC-NER assay in NPE that was supplemented with DMSO or 200 µM MLN4924, which prevents cullin neddylation. See Figure S2K for western blot of extracts used. **(G)** Schematic illustrating the gap filling assay. Staggered cutting by EcoRI and KpnI (orange lines) allowed resolution of the top (non-template) and bottom (template) strands. **(H)** Unscheduled DNA Synthesis (UDS) assay. Repair reactions were performed as in Figure 1D, except that GG-NER was inhibited by immunodepletion of XPC from NPE, and that NPE was supplemented with [α-^32^P]dCTP. DNA was recovered, digested with EcoRI and KpnI to distinguish top and bottom strands (G), separated on a Urea-PAGE gel, and subjected to autoradiography.

### Structure of a TC-NER complex containing STK19

To determine how STK19 promotes repair, we used AlphaFold-Multimer (AF-M) to screen for potential STK19 interactors. STK19 was “folded” with 409 proteins involved in genome maintenance and transcription, and each binary structure prediction was assessed using SPOC (Structure Prediction and Omics Classifier), a classifier trained to distinguish functionally relevant from spurious AF-M complexes (0-1 scale; >0.5 is a strong candidate for a meaningful interaction).^27^ Among the proteins examined, RPB1 (Pol II subunit 1), XPD (ERCC2), and CSA (ERCC8) were the top hits (Figure 3A; see Figures S3A and S3B for conventional confidence metrics; Table S1). By folding CSB, CSA, DDB1, DDA1 (a component of CRL4 E3 ligases),^28^ ELOF1, UVSSA, and STK19 all at once, we generated a structure prediction for an STK19-containing TC-NER complex (Figures 3B and 3C; but lacking Pol II) that allowed us to initiate structure-function analyses of STK19 and other TC-NER proteins (see below).

**Figure 3.**
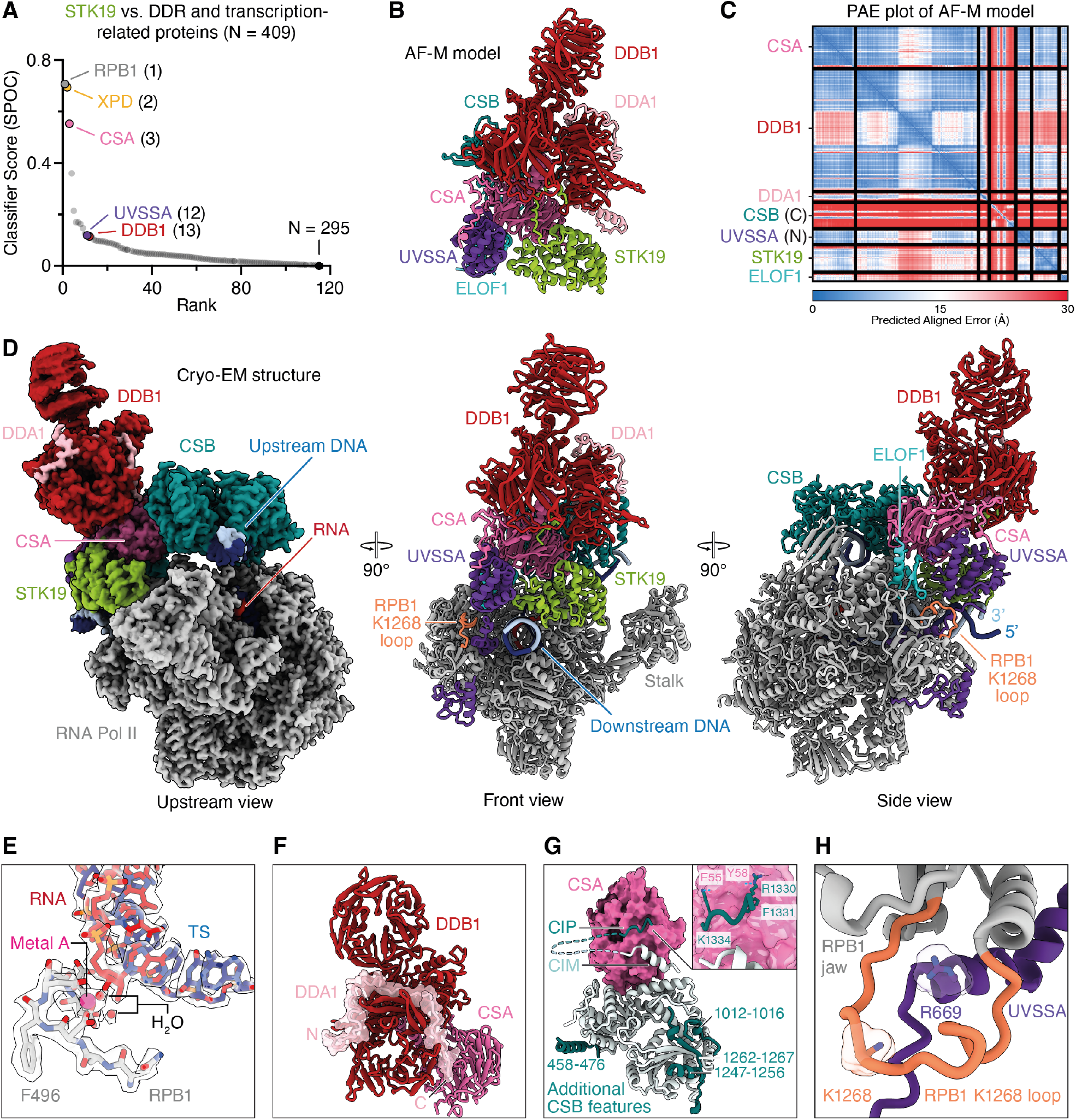
Structure of an STK19-containing TC-NER complex. **(A)** STK19 was “folded” with ∼400 proteins, and the resulting structure predictions were ranked by SPOC, a classifier trained to distinguish true from spurious AF-M predictions. **(B)** Human CSA, DDB1, DDA1, CSB (aa 1250-1493), UVSSA (aa 1-150), STK19, and ELOF1 were folded in five models using AF-M, and a representative structure prediction is shown. The structure has confidence metrics: pLDDT = 82.7, PAE = 3.8, avg_models = 0.86. **(C)** Representative predicted alignment error (PAE) plot for the structure in (B). **(D)** Three views of the STK19-containing TC-NER complex by cryo-EM. Structure is presented as a cartoon model (front and side views) or as the Coulomb potential (map xi). Template strand, dark blue. Non-template strand, light blue. DDB1 β-propeller 2 is shown as low-pass filtered map (map i) superposed on map ix. **(E)** Stick representation of Pol II active site. Coulomb potential map is shown as transparent volume. Metal A and two coordinating water molecules are clearly visible and shown as pink or red spheres, respectively. **(F)** Interaction of DDB1, CSA, and DDA1. DDA1 is shown in surface and cartoon representation. **(G)** New CSB features and interaction of CSA and CSB. CSA shown as surface, and CSB shown as cartoon model. Additional CSB features are colored in dark cyan. An N-terminal CSB α-helix (residues 458-476) binds to ATPase lobe 1, and a β-strand (residues 1262-1267) complements the β-sheet of ATPase lobe 2 in an anti-parallel orientation. The additional CSB β-strand is flanked by two α-helical elements (CSB residues 1012-1016 and 1247-1256). Inset, details of CSB’s newly resolved CSA-interacting peptide (CIP) shown in cartoon representation with important side chains shown in stick representation. **(H)** RPB1 K1268 loop shown with C-terminus of UVSSA. RPB1 K1268 contacts the start of the UVSSA C-terminus. UVSSA R669 inserts into the loop. UVSSA R669 and RPB1 K1268 residues are shown in stick and surface representation.

We subsequently used single-particle cryo-EM to solve the structure of STK19 bound to a Pol II elongation complex containing CSB, CSA-DDB1, DDA1, ELOF1, and UVSSA (Figure S4). We collected and analyzed a cryo-EM dataset (Figures S5 and S6) and obtained a structure at an overall resolution of 1.9 Å from 484,012 particles (Figure 3D). In our structure, Pol II adopts a post-translocated state, and we resolve water molecules that coordinate the metal A in the active site (Figure 3E). CSB embraces the upstream DNA, and its ATPase motor is in a pre-translocated state.^11^ The atomic model of the TC-NER complex was real-space refined and shows excellent stereochemistry (Tables S2 and S3). High-resolution features allowed us to unambiguously place a structure of the Pol II-TC-NER complex with ELOF1 into our density.^15^

We also observed well-resolved features for STK19, allowing us to unambiguously dock an AlphaFold model of STK19 into the corresponding density, which showed that STK19 binds the TC-NER complex primarily via RPB1 and an extensive interface with CSA (Figure 3D; see below for a detailed description), as predicted by AF-M (Figure 3B). Additional density corresponding to DDA1 on DDB1 could be built using an AF-M prediction. DDA1 interacts with DDB1 as observed before^29^ (Figure 3F).

Our TC-NER complex also contained additional densities, leading to a more complete model of this assembly. First, we modeled additional N- and C-terminal parts of CSB that bind to ATPase lobes 1 and 2, respectively (Figure 3G). Second, besides the known interaction of CSA with the CIM of CSB,^11,12^ we also observed a contact between highly conserved CSB residues 1329-1336 and CSA, which we name CSA-interacting peptide (CIP) (Figures 3G and S3C). Specifically, CSB R1330 contacts CSA Y58, F1331 of CSB inserts into a cavity formed between WD40 repeats 1 and 2 of CSA, and CSB K1334 forms a salt bridge with E55 of CSA. Third, we extended the model for the linker region between the UVSSA zinc finger domain and the UVSSA C-terminus and completed the DDB1 model by resolving additional residues in the previously unobserved flexible loops. Fourth, the previously unresolved C-terminal tail of CSA was seen to interact with DDB1 and the VHS domain of UVSSA (Figure 3D; described in more detail below). Lastly, we resolved and modeled an additional loop of the RPB1 jaw (residues 1261-1281) containing K1268, which is ubiquitinated during TC-NER.^16,30^ Our structure shows that the loop is positioned above the UVSSA C-terminus, with UVSSA residue R669 inserting into the cavity formed by the RPB1 jaw and the K1268 loop (Figures 3D and 3H). Remarkably, almost all of the above features were correctly predicted by AF-M (Figure 3B; Figures S3D and S3E; see below).

### The interaction of STK19 with CSA and DDB1 is required for TC-NER

STK19 comprises three winged helix (WH) domains^18,19^ (Figures 4A and 4B) and it contacts the TC-NER complex in four places (Figure 4C). First, STK19 density could be traced from WH1 towards a pocket formed by DDB1 β-propellers A and B (Figure 4D, panel I). Directed by AF-M predictions (Figure 3B), we assigned this additional density to the previously unresolved N-terminus of STK19, which binds to the same DDB1 cavity that is also occupied by the N-terminus of CSA. Second, STK19 residues R72, T73, D76, and R77 in the α3 helix of WH1 contact the linker between CSA WD40 repeats 5 and 6 (Figure 4D, panel II). STK19 destabilizes a CSA loop (residues 231-236) normally seen between WD40 repeats 4 and 5^15^ that we could no longer resolve fully. Third, WH2 and the N-terminal part of the WH3 α1 helix directly interact with the RPB1 clamp head (Figure 4D, panel III). Finally, STK19 WH3 inserts into a pocket formed by downstream DNA, UVSSA, and CSA, where K203 in WH3 contacts E10 in UVSSA (Figures 3D and 4D, panel IV). Compared to the previous TC-NER complex with ELOF1,^15^ the UVSSA VHS domain is shifted towards the downstream DNA and STK19 by 5-8 Å (Figure 4E). Of note, the only experimentally observed interaction among TC-NER factors not confidently predicted by AF-M in a pairwise “folding” experiment was the one between UVSSA and STK19. Consistently, UVSSA^E10A^ supported efficient TC-NER (Figure 4F), suggesting that the STK19-UVSSA interaction is predominantly facilitated by their common binding partner CSA (Figures 3B and 3D). To further address whether binding of STK19 to the TC-NER complex is important for repair, we deleted the N-terminus of STK19, which projects towards DDB1 (Figure 4D, panel I), resulting in STK19^DN^, or we mutated the four residues at the STK19-CSA interface to alanine (Figure 4D, panel II; *X. laevis* residues in parentheses), yielding STK19^4A^. Both of these mutants supported only ∼50% efficient TC-NER when added to STK19-depleted extract, and combining these mutations lead to a complete loss of repair (Figure 4G). Similarly, reversing two charges at the CSA-STK19 interface strongly impaired error-free repair (Figure 4G; STK19^DR-RD^). We conclude that the interaction of STK19 with the TC-NER complex via interfaces with DDB1 and CSA is essential for STK19 to support cell-free repair.

**Figure 4.**
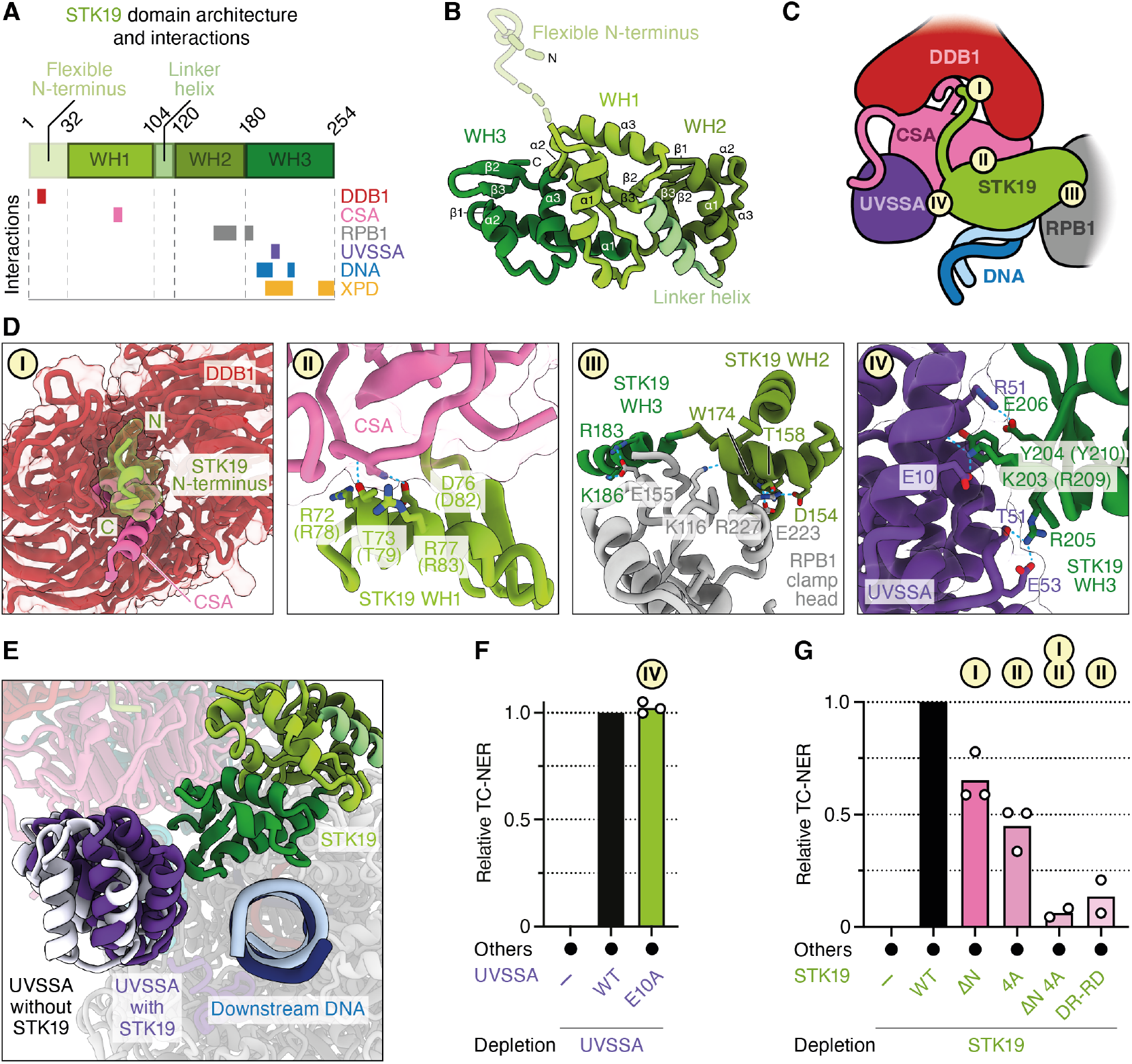
STK19 interacts with the TC-NER complex via CSA and DDB1. **(A)** Schematic showing the domain architecture of STK19 and interaction partners for each subdomain. **(B)** Cryo-EM structure of STK19 colored by domain architecture. Unresolved residues are shown as dotted lines. **(C)** Schematic showing STK19 and its interaction partners, including DNA. **(D)** Close-up view of interaction sites I-IV from (C). STK19, UVSSA, and DDB1 are shown in cartoon and/or surface representation. CSA is shown as a cartoon model. STK19 N-terminal backbone could be resolved, amino acid register is not assigned. Residues forming hydrogen bonds at the interaction interfaces are shown in stick representation. Hydrogen bonds shown as dotted lines. *X. laevis* residues are shown in parentheses. **(E)** Binding of STK19 to the TC-NER complex induces a shift of the UVSSA VHS domain towards the downstream DNA and STK19 by up to 8 Å. UVSSA VHS domain from TC-NER complex without STK19 shown in white (PDB ID 8B3D). **(F)** TC-NER assay in UVSSA-depleted NPE. *X. laevis* UVSSA proteins expressed in wheat germ extract were added as indicated. **(G)** TC-NER assay in STK19-depleted NPE. Buffer or the indicated *X. laevis* STK19 variants (Figure S2B) were added back. ΔN, deletion of amino acids 1-33; 4A, mutation of residues R78, T79, D82, and R83 to alanine; ΔN 4A, mutant containing both ΔN and 4A mutations; DR-RD, residues D82 and R83 were swapped.

### Mutations at the predicted STK19-XPD interface disrupt TC-NER

While our results show that STK19 binding to CSA and DDB1 is important for TC-NER, they do not explain how this binding promotes repair. Importantly, our *in silico* screen predicted that STK19 also interacts with the XPD subunit of TFIIH via an extensive interface involving STK19’s WH3 domain (Figure 5A). This interaction has a high SPOC score in humans (Figure 3A), and it is confidently predicted in other species that have STK19 and XPD, including frogs, fish, plants, and fission yeast (Figure S3F). Based on these structure predictions, we postulated that STK19 positions TFIIH on the TC-NER complex. Of note, the XPD and UVSSA binding sites on STK19 partly overlap (Figures 5A and 5B), and we provide evidence below suggesting that UVSSA moves to allow formation of the XPD-STK19 complex (Figure 5C and see below). To address the importance of the STK19-XPD interaction, we generated three STK19 mutants that are designed to disrupt three distinct contact points between STK19 and XPD (Figure 5D, panels I-III). These mutations involved changing charged and bulky residues in a loop to two glycines (STK19^RY-GG^; R209G and Y210G in *X. laevis*), a single charge-swap mutation (STK19^D-R^; D217R), and replacement of two small side chains in a loop to two larger residues (STK19^AS-EY^; A250E and S251Y). These mutations reduced TC-NER to various extents, and when all the mutations above were combined, repair was inhibited up to ∼4 fold (Figure 5E). Although the STK19^RY-GG^ mutant also removes a salt bridge with E10 of UVSSA (Figure 4D, panel IV), we showed above that UVSSA^E10A^ is fully proficient for repair. Therefore, the effect of this mutant is likely due to deficient XPD binding. Together, our data suggest that STK19 promotes TC-NER by interacting with XPD.

**Figure 5.**
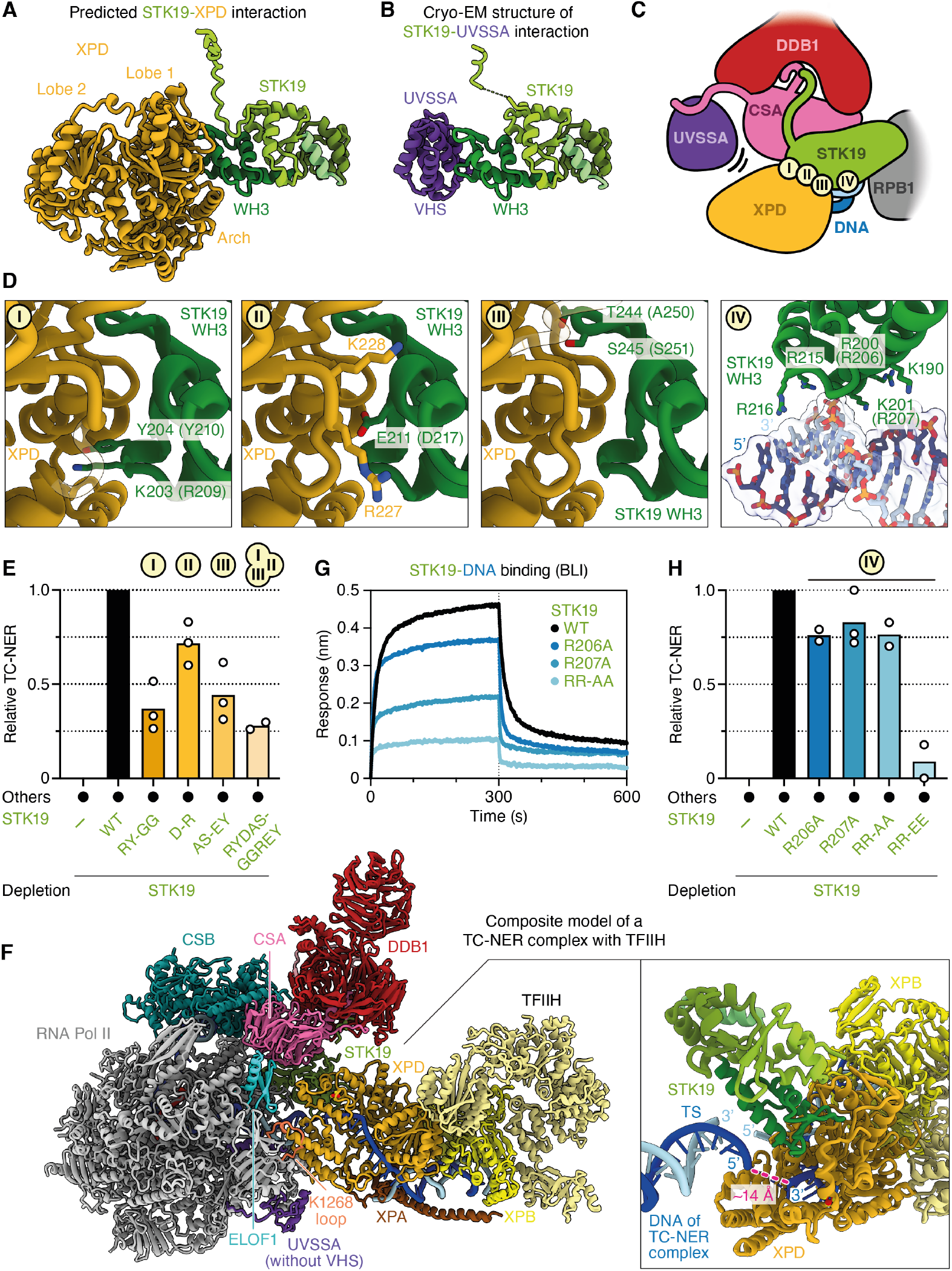
Mutations at the predicted STK19-XPD interface disrupt TC-NER, but STK19 DNA binding is not required. **(A-B)** AF-M prediction for STK19-XPD complex (A) and cryo-EM structure of STK19-UVSSA shown in Figure 3D (B). STK19 is depicted in the same orientation in both panels. **(C)** Schematic showing STK19 and its interaction partners, including XPD. The proposed dissociation of UVSSA from CSA’s β-propeller that allows XPD binding to STK19 is indicated. **(D)** Close up view of interaction sites I-IV from (C). Amino acids shown in parentheses refer to *X. laevis*. **(E)** TC-NER assay in STK19-depleted NPE. Buffer or the indicated *X. laevis* STK19 mutants (Figure S2B) were added back. RY-GG, mutation of R209 and Y210 within a loop in WH3 to glycine; D-R, D217R; AS-EY, residues A250 (corresponding to T244 in *H. sapiens*) and S251 located in a loop within WH3 were mutated to larger residues to interfere with XPD recruitment. **(F)** Model for the positioning of TFIIH on the TC-NER complex mediated by STK19. The STK19-XPD structure prediction was aligned with STK19 in the TC-NER complex shown in Figure 3D. Subsequently, a TFIIH-XPA-DNA complex (PDB ID 6RO4) was aligned with XPD from the STK19-XPD structure prediction. The UVSSA VHS domain and CSA’s C-terminal tail were removed for clarity and to reflect our model that the VHS domain moves to accommodate XPD. Inset, close-up of the STK19-XPD-DNA region. The distance between the template strand of the TC-NER complex and the ssDNA in XPD of the TFIIH complex is shown. **(G)** Biolayer interferometry (BLI) assay measuring the interaction of *X. laevis* STK19 WT and the indicated mutants (Figure S2B) with a biotinylated 14 nt DNA duplex. RR-AA, combination of R206A and R207A. **(H)** TC-NER assay in STK19-depleted NPE. Buffer or *X. laevis* STK19 mutants described in (G) were added. RR-EE, charge-swap mutation of R206 and R207 to glutamate.

### A model of the TC-NER complex in which STK19 positions XPD on DNA

XPD is the 5’ to 3’ helicase subunit of TFIIH that tracks along the transcribed strand during TC-NER to verify the presence of damage. We asked how the interaction of STK19 with XPD would position TFIIH relative to the TC-NER complex. To this end, we first aligned the STK19-XPD AF-M prediction (Figure 5A) on the cryo-EM structure of the TC-NER complex via STK19 (Figure 5F). Onto the resulting complex, we aligned a previously determined TFIIH-XPA-splayed DNA structure^31^ via XPD (Figure 5F). This revealed no major clashes between TFIIH and the TC-NER complex other than the one between XPD and UVSSA mentioned above. In the composite model, STK19 positions TFIIH in front of the TC-NER complex near the downstream DNA, where it searches the template strand for the presence of DNA damage. Strikingly, the 3’ end of the single-stranded DNA emerging from the XPD channel aligns with the 5’ end of the downstream template strand of the TC-NER complex (Figure 5F, right panel, dark blue strands), suggesting that STK19 guides TFIIH to the correct strand for lesion verification. Importantly, in the TC-NER cryo-EM structure, STK19 also contacts downstream DNA (Figure 5D, panel IV), consistent with prior DNA binding studies.^18,19^ Specifically, a positively charged surface in the WH3 domain contacts the phosphate backbone. To address the importance of this interaction, we mutated two arginines to alanine (R206A and R207A in *X. laevis*), alone or in combination. The single mutants reduced and the double mutant (STK19^RR-AA^) strongly impaired DNA binding (Figure 5G). Despite this, all three mutants including STK19^RR-AA^ had only a minor effect on DNA repair (Figure 5H). In contrast, a mutant in which both charges were reversed (STK19^RR-EE^) was severely defective in DNA repair. These results suggest that STK19 does not need to attract DNA, but that repelling it is deleterious. Indeed, as shown in Figure 5F (right panel), duplex DNA contacts STK19 immediately adjacent to the place where the template strand enters the XPD channel. Therefore, we propose that STK19 must accommodate DNA in this location to allow template strand entry to the XPD channel.

### A novel interaction between CSA and UVSSA

Our model above proposes that STK19 positions TFIIH via XPD binding, yet XPD’s position on STK19 would clash with the VHS domain of UVSSA, which is anchored nearby via CSA (compare Figures 5C and 6A). Importantly, the VHS domain interacts with CSA in two ways (Figures 6A and 6B, panels I and II): the previously reported interaction with the CSA β-propeller involving Y334 (Figure 6B, panel I) and a novel interaction involving the C-terminal tail of CSA (panel II), revealed by AF-M and cryo-EM (Figures 3B and 3D). Several proximal residues in this ∼30 amino acid long tail interact with DDB1 (Figure 6B, panel III), and the distal W389 makes a contact with the UVSSA VHS domain (Figure 6B, panel II) that is highly conserved (Figures S7A and S7B). To determine which of these contacts are important to mediate UVSSA-CRL4^CSA^ binding, we measured CRL4^CSA^-dependent UVSSA ubiquitination *in vitro*. As shown in Figure 6C, mutation of the tryptophan and the adjacent, conserved serine specifically prevented UVSSA monoubiquitination by CRL4^CSA^ (Figures 6C and S7C; STK19^WS-AA^). These results indicate that CSA’s C-terminus is required to mediate a stable interaction between CRL4^CSA^ and UVSSA. Importantly, these mutations also abolished TC-NER (Figure 6D). In contrast, mutation of the tyrosine that resides at the previously described CSA-UVSSA interface (Figure 6B, panel I) had no effect on UVSSA ubiquitination (Figure 6C; STK19^Y-A^) or TC-NER (Figure 6D). These results suggest that binding of CSA’s flexible C-terminal tail to UVSSA is more important than binding mediated by CSA’s β-propeller. We therefore propose that UVSSA can dissociate from the CSA β-propeller to accommodate XPD while remaining tethered to the TC-NER complex via CSA’s C-terminal tail (Figure 5C). Together, our results suggest a model that explains how STK19 functionally couples the TC-NER complex to downstream repair events (Figure 7 and see Discussion).

**Figure 6.**
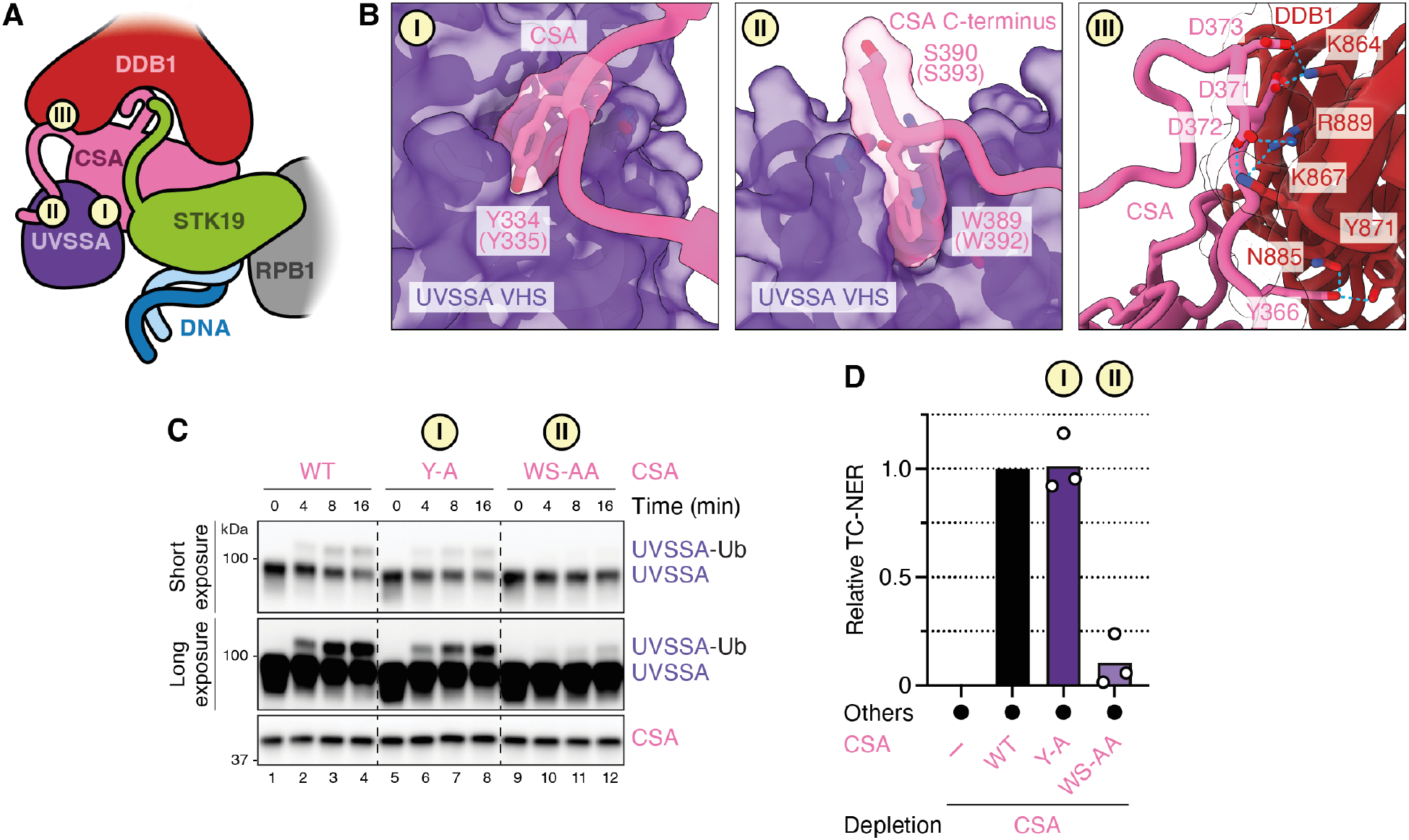
The C-terminus of CSA binds UVSSA and is essential for TC-NER. **(A)** Schematic showing that the VHS domain of UVSSA binds CSA at two points: (I) the CSA β-propeller and (II) the CSA C-terminal flexible tail (distal region). (III) Interaction of CSA’s C-terminal tail (proximal region) with DDB1. **(B)** Close up view of interaction sites I-III from (A). Amino acids shown in parentheses refer to *X. laevis*. **(C)** *In vitro* ubiquitination assay. *X. laevis* UVSSA was mixed with neddylated *X. laevis* CRL4^CSA^ variants (WT, Y-A, or WS-AA; Figure S2B), ubiquitin, E1, UBE2E1, and ATP. At the indicated times, reaction products were blotted for UVSSA and CSA. UVSSA-Ub, monoubiquitinated UVSSA. Y-A, *X. laevis* Y335 mutated to alanine. WS-AA, *X. laevis* W392 and S393 mutated to alanine. **(D)** TC-NER assay in CSA-depleted NPE. Buffer or the indicated *X. laevis* CRL4^CSA^ mutants described in (C) were added back.

**Figure 7.**
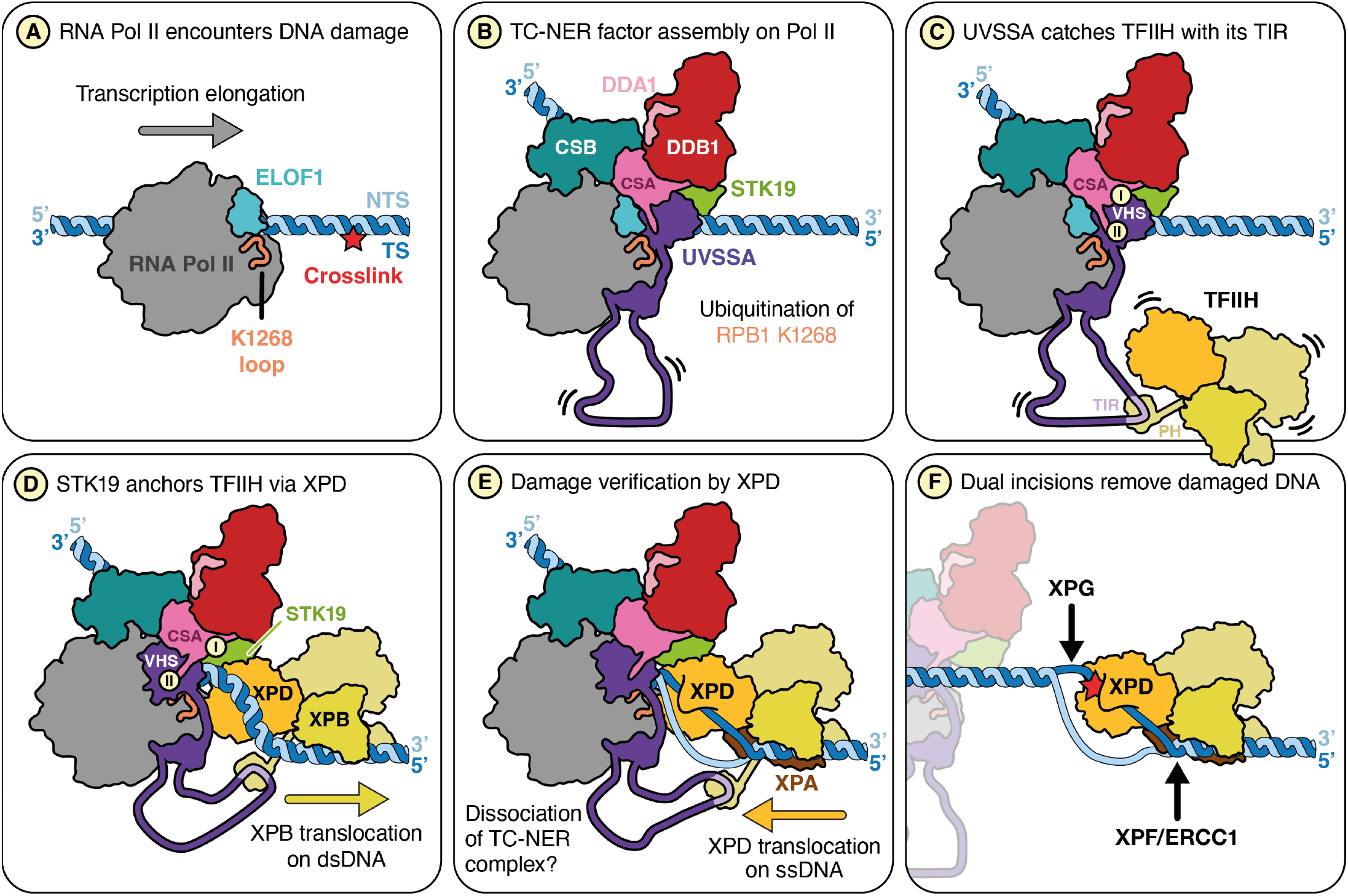
Model of STK19 function in TC-NER. **(A)** Pol II stalls at a lesion (red star) in the template strand. **(B)** TC-NER factors (CSB, CRL4^CSA^, UVSSA, ELOF1, and STK19) are assembled on stalled Pol II. Lysine residue 1268 of RPB1 (part of orange loop) is ubiquitinated by CRL4^CSA^, which docks onto ELOF1 (ubiquitin not shown). **(C)** The TFIIH-interacting region (TIR) of UVSSA binds to the p62 subunit of TFIIH, tethering TFIIH to the TC-NER complex. Since the TIR is located within a long, unstructured region of UVSSA (shown as a long loop), TFIIH can access a large area around the TC-NER complex. In this state, the N-terminal VHS domain of UVSSA interacts with the CSA β-propeller (I) as well as CSA’s C-terminal tail (II). **(D)** TFIIH is positioned by STK19 close to the downstream DNA. The VHS domain of UVSSA dissociates from the CSA β-propeller (I) but remains associated with CSA’s C-terminus (II), allowing XPD to bind STK19. Due to STK19’s interaction with XPD, TFIIH and therefore XPB are anchored relative to the damaged DNA. As a result, XPB translocation away from the TC-NER complex pumps DNA into the space between XPB and STK19, leading to DNA unwinding and template strand association with the XPD helicase channel. **(E)** XPD translocates along the template strand in the 5’–3’ direction to verify the damage. It is unclear whether the TC-NER complex is pushed back by TFIIH or whether it dissociates from DNA. This intermediate most closely resembles the structure modeled in Figure 5E. **(F)** The structure-specific endonucleases XPF-ERCC1 and XPG perform dual incisions to remove the damaged DNA. Subsequent gap filling (not depicted) completes repair.

## DISCUSSION

We have recapitulated transcription-coupled DNA repair in frog egg extracts, and we use this system to show that STK19 is the missing link between stalled Pol II and TFIIH. We find that extracts depleted of STK19 are deficient for TC-NER as seen from failure to restore a restriction site at a cisplatin lesion in the template strand. Together with mutational analyses, our 1.9 Å resolution cryo-EM structure of the STK19-containing TC-NER complex demonstrates that the interaction of STK19 with CSA-DDB1 is critical for repair. Structure prediction-guided mutagenesis further indicates that STK19 also interacts functionally with the XPD subunit of TFIIH. Finally, we identify a novel interface between the C-terminal tail of CSA and UVSSA that is essential for TC-NER. Together, our results suggest a model of how STK19 promotes TC-NER (Figure 7).

### STK19: lynchpin between TFIIH and stalled Pol II

Previous studies showed that after CSB binds to stalled Pol II, CRL4^CSA^ and UVSSA are recruited and collaborate with ELOF1 to promote Pol II ubiquitination and subsequent TFIIH recruitment (Figures 7A and 7B).^9^ Furthermore, UVSSA was shown to bind TFIIH to the TC-NER complex via its TFIIH-interacting region (TIR) that we confirm is essential in cell-free TC-NER (Figure 7C and Figure 2E). Importantly, because the TIR motif is located in an unstructured part of UVSSA (Figure S7D), the tethered TFIIH has many degrees of freedom (Figure 7C), and it was unclear how it is positioned on the template strand ahead of Pol II for lesion verification. We propose that after being tethered via UVSSA (Figure 7C), TFIIH docks onto STK19 via XPD, which guides TFIIH to the DNA downstream of stalled Pol II (Figure 7D). Subsequently, XPB translocates away from stalled Pol II by tracking along the template strand in the 3’–5’ direction, which unwinds DNA, allowing XPD to capture ssDNA in its helicase channel (Figure 7E).^3^ Indeed, in the full TC-NER-TFIIH model, the damaged strand in TFIIH is located only ∼14 Å from the transcribed strand in Pol II (Figure 5F). Because a charge-swap but not an alanine substitution in the STK19 DNA-binding site impairs TC-NER (Figure 5H), we propose that STK19 must accommodate downstream DNA near the XPD helicase channel to guide the template strand into the ATPase. XPD then translocates in the 5’–3’ direction, searching for DNA damage (Figure 7E). When a lesion is located, XPD stalls, followed possibly by further DNA unwinding by XPB,^32^ dual incisions by XPF-ERCC1 and XPG (Figure 7F), and gap filling (not depicted).

In our model, XPD binding to STK19 would overlap with UVSSA’s VHS domain in the TC-NER complex (Figures 7C and 7D; Figures 5A and 5B). Importantly, in addition to the previously reported contact between UVSSA and the β-propeller of CSA (Figure 7C, label I), UVSSA also binds the flexible C-terminal tail of CSA (Figure 7C, label II). Our mutagenesis results further suggest that the latter interaction is more important than the former. Based on these results, we propose that UVSSA dissociation from the CSA β-propeller makes room for XPD to bind STK19, even as UVSSA remains attached to the CSA C-terminus (Figure 7D, label II). In this way, UVSSA first tethers TFIIH to the TC-NER complex and then delivers it to STK19 without allowing TFIIH dissociation.

Interestingly, STK19 depletion reduced repair to a greater extent than the combination of five point mutations targeting the STK19-XPD interface. One explanation of this difference is that these mutations did not fully disrupt the STK19-XPD interaction. Alternatively, STK19 may induce allosteric changes in the TC-NER complex to enhance repair. For example, in the presence of STK19, the UVSSA VHS domain moves towards DNA by 5-8 Å, which might enhance its affinity for the TC-NER complex. In this view, even without a direct XPD-STK19 interaction, TFIIH can eventually find its proper location on the DNA, albeit inefficiently.

### Common themes in transcription initiation, GG-NER, and TC-NER

TFIIH is essential for DNA opening during transcription initiation, GG-NER, and TC-NER.^3^ Based on our findings and those of others, we propose that TFIIH function in these three pathways involves at least two common principles. First, the initial recruitment of TFIIH in all three pathways occurs via the PH domain of the p62 subunit. Thus, acidic sequences in the general transcription factor TFIIE, XPC, and UVSSA bind to the same basic groove in the PH domain with high affinity.^17^ The second common principle involves the mechanism of DNA opening by the XPB ATPase subunit of TFIIH. During transcription initiation, XPB translocates along duplex DNA while TFIIH is also anchored to the Pol II stalk.^33^ This configuration effectively pumps DNA into a gap between the two TFIIH contact points, leading to DNA unwinding. Similarly, in GG-NER, TFIIH is attached via multiple contacts to XPC, which grips the damaged DNA at a distance from XPB.^32^ Finally, our data indicate that in the case of TC-NER, TFIIH anchoring involves XPD docking onto the TC-NER complex via STK19 so that XPB pumping DNA towards the stalled polymerase leads to strand separation (Figures 7D and 5F). These considerations argue that both TFIIH recruitment and anchoring are generally required for its function in transcription initiation and repair. Notably, a key difference between these processes is that unlike in transcription initiation, GG-NER and TC-NER require the ATPase activity of XPD,^3^ and XPD is positioned immediately adjacent to the DNA (Figure 5D and ^32^). As a result, once the duplex is unwound by XPB, one strand readily enters the XPD helicase channel for lesion verification.

### A possible function for K1268 ubiquitination in TC-NER

The ubiquitination of RPB1 K1268 stimulates the association of TFIIH with the TC-NER complex, but the underlying mechanism is not understood.^16,30^ Our high resolution cryo-EM structure allowed us to model RPB1’s K1268 loop. The loop is positioned such that residue K1268 points towards the base of UVSSA’s C-terminal helix and the VHS domain (Figure 3H). It is therefore tempting to speculate that RPB1 ubiquitination remodels the loop and surrounding features, and that this promotes the UVSSA VHS domain’s repositioning that we propose is essential for XPD binding to STK19. Strikingly, the K1268 loop is also located at the interface with XPD in the composite TC-NER model containing TFIIH (Figure 5F).

Our model of TFIIH positioning, together with structural information about K1268’s location, sets the stage to address how RPB1 ubiquitination promotes TC-NER.

### The power of *in silico* screening for protein-protein interactions

To understand how STK19 functions, we initially used AlphaFold-Multimer (AF-M) to screen for STK19 partners among ∼400 proteins involved in genome maintenance and transcription. Given this limited search space, conventional confidence metrics were adequate to identify CSA, XPD, and RPB1 as top candidates. However, SPOC, a classifier trained to identify functionally relevant structure predictions, gave higher relative scores to STK19’s functional partners than conventional metrics, consistent with its greater discriminatory power (Figure 3A vs. Figures S3A and S3B). These predictions immediately suggested a hypothesis for STK19’s mechanism and allowed us to engineer site-directed mutants that test the model. Subsequently, we determined the cryo-EM structure of the STK19-containing TC-NER complex (Figure 3D), which extended and provided critical experimental support for the structure predictions. This example further illustrates the remarkable synergy between structure prediction and experimental structure determination that accelerates mechanistic discovery.

### A cell-free system for TC-NER

We report the first cell-free system that recapitulates all key features of TC-NER. Repair in this system requires transcription, all known TC-NER factors, interactions among these factors, and CRL E3 ligase activity, and it is accompanied by gap filling in the template strand. Our approach is ideally suited to address key questions in the field such as the role of ubiquitination and the mechanism of transcription resumption after repair is complete.

## ACKNOWLEDGEMENTS

We are grateful to Stephen Buratowski and members of his group for providing numerous reagents and for many helpful discussions, which were instrumental in helping us develop cell-free transcription. We thank Daniel Durocher and Dheva Setiaputra for alerting us to the predicted interaction between STK19 and CSA. Thanks to Seychelle Vos, Karen Adelman, Maria Jessica Bruzzone and members of the Walter laboratory for feedback on the manuscript. We thank the Harvard Cryo-EM Center for Structural Biology at Harvard Medical School for support with data collection. T.E.T.M. was supported by an HHMI fellowship of The Jane Coffin Childs Memorial Fund for Medical Research and an EMBO Long-term fellowship (ALTF 1316-2016). L.F. is supported by the NIH Director’s New Innovator Award (DP2-ES036404). This work was supported by NIH grant HL098316 to J.C.W., who is a Howard Hughes Medical Institute Investigator and an American Cancer Society Research Professor.

## AUTHOR CONTRIBUTIONS

J.C.W and T.E.T.M. conceived and T.E.T.M. performed and analyzed all experiments except for cryo-EM studies. T.E.T.M carried out structure predictions, and SPOC analysis was performed by E.W.S. Cryo-EM protein preparation, complex formation, and cryo-EM analysis were conceived and performed by M.K. and L.F. The atomic model was built by L.F. The manuscript was written by T.E.T.M and J.C.W., with help from L.F.

## METHODS

### Preparation of DNA substrates

All DNA plasmids used for *in vitro* transcription and TC-NER in this study are listed in Table S4. A plasmid for *in vitro* transcription (pUC18-G5AdML(Δ53)G-) was a generous gift of Stephen Buratowski, and was modified by “Round-the-Horn” PCR to place a tandem BbsI restriction cassette downstream of the promoter for the subsequent introduction of a site-specific lesion. The resulting plasmid pTM07_5xUAS_AdML^Δ53^_BbsI, which we refer to as pAdML^Δ53^, contains five Upstream Activating Sequences (UAS), a truncated adenovirus major late promoter directly upstream of a G-less cassette, and the BbsI cassette. To increase the transcriptional output, we replaced the AdML^Δ53^ as well as the adjacent G-less cassette with the super core promoter 2 (SCP2)^34^ and a fragment of the *X. laevis* ubiquitin gene. We subsequently changed the SCP2 Initiator from 5’-TCAGAC-3’ to 5’-TCAGTC-3’ to maximize the use of a single transcription start site (TSS). We refer to this modified SCP2 promoter as SCP2*. The resulting second generation IVT plasmid (pTM171_ 5xUAS_SCP2*_Ub_BbsI; referred to as pSCP2*) was assembled from multiple gene blocks ordered from Integrated DNA Technologies (IDT) using Gibson Assembly (NEBuilder^®^ HiFi DNA Assembly Master Mix, New England Biolabs, #E2621L). Additionally, we introduced polyA sites upstream of the 5xUAS to prevent multiple rounds of transcription, and altered the size of the ubiquitin gene to position the BbsI cassette at different locations downstream of the TSS. These modifications of pSCP2* generated pCtrl-122 and pCtrl-322 (Table S4). Other plasmids used for IVT reactions were ordered from Addgene (pCMV; cat #11153) or assembled as described previously (pActin).^25^

The generation of plasmids containing a site-specific cisplatin 1,3-GTG intrastrand crosslink involved the following steps. First, pCtrl-122 and pCtrl-322 were digested with BbsI-HF (NEB, R3539L) and purified on a HiLoad 16/600 Superdex 200 pg column (Cytiva). Second, the preparation of the lesion-containing insert was performed as described previously.^35^ In short, a DNA oligo containing a unique GTG site (5’–CCC TCT CCA CGT GTC TCC TC-3’) was platinated and purified on a Mono Q 5/50 GL column (Cytiva) before it was annealed to the complementary strand (5’–GCA CGA GGA GAC ACG TGG AG–3’). Third, the lesion-containing duplex DNA was ligated into the purified, linear backbones of pCtrl-122 and pCtrl-322. Subsequent purification was performed using CsCl gradient centrifugation and butanol extraction. The final lesion-containing plasmids pPt-122 and pPt-322 were verified by PmlI restriction digestion, indicating that more than 95% of the plasmids contained the crosslink. Lesion plasmids were frozen in liquid nitrogen and stored at −80°C.

### *In vitro* transcription (IVT) assay

Xenopus egg extracts (HSS, high-speed supernatant; NPE, nucleoplasmic extract) were prepared as described.^36^ Thawed egg extracts were supplemented with ATP regenerating system (2 mM ATP, 20 mM phosphocreatine, and 5 μg/ml phosphorkinase; final concentrations), 2 mM DTT (only NPE), and 3 µg/ml nocodazole (only HSS).

HSS was cleared by centrifugation at 14,000 x g for 5 min at room temperature prior to IVT reactions. Where indicated, recombinant transcription activator GAL4-VP64 (1 µM final concentration in IVT reaction), recombinant human TBP (500 nM), and α-amanitin (2 µM) were added to egg extracts. The resulting “Extract Mixture” was incubated for 15 min at room temperature. In parallel, an “IVT Mixture” was prepared with the follow-ing components: 10x IVT Buffer–100 (200 mM HEPES pH 7.5, 1 M KCl, 50 mM MgCl_2_, and 5 mM EDTA; 1x final concentration in IVT reaction), Recombinant RNasin™ Ribonuclease Inhibitor (Promega; 0.5 U/µl), indicated IVT plasmid (7 ng/µl), [α-^32^P]UTP (Revvity; 0.5 µCi/µl), and, unless otherwise indicated, polyethylene glycol 20,000 (PEG^20K^; Hampton Research; 1% (v/v)). To initiate IVT reactions, “Extract Mixture” and “IVT

Mixture” were combined such that egg extracts (supplemented with ATP regenerating system and DTT or nocodazole) represent 50% of the final reaction volume. Reactions were incubated at room temperature. At indicated times, samples were withdrawn and mixed with seven volumes of IVT Stop Solution (50 mM Tris pH 8.0, 25 mM EDTA, and 0.5% SDS) and treated with 1.2 mg/ml Proteinase K (Roche) for 1 h at 37ºC. RNA transcripts were then purified using RNAClean XP (Beckman), according to manufacturer’s instructions, except that two volumes of bead solution were added and RNA was eluted with TE Buffer. Eluted RNA transcripts were mixed with 2x IVT Loading Buffer (100% formamide supplemented with 1 mM EDTA, 0.1% SDS, 1 µg/ml Xylene Cyanol FF, and 1 µg/ml Bromophenol Blue), heated to 95ºC for 5 min, rapidly cooled on ice, and resolved on an 8% Urea-PAGE gel in 0.8x Glycerol Tolerant Gel Buffer (20x stock: 1.78 M Tris, 0.57 M taurine, 0.01 M EDTA). Gels were dried under vacuum, exposed to a phosphor screen, and imaged on a Typhoon™ FLA 7000 phosphorimager (GE Healthcare). RNA transcripts in Figures 1B and S1B were purified by conventional phenol-chloroform extraction and ethanol precipitation instead of using RNAClean XP beads.

### Cell-free TC-NER assay

The repair of plasmids containing a cisplatin 1,3-GTG intrastrand crosslink was performed under IVT assay conditions, with the following changes. The “IVT Mixture” contained 10x IVT Buffer–33 (200 mM HEPES pH 7.5, 330 mM KCl, 50 Mm MgCl_2_, and 5 mM EDTA; 1x final concentration in IVT reaction) instead of 10x IVT Buffer–100, damaged IVT plasmid (10 ng/µl instead of 7 ng/µl), and no [α-^32^P]UTP. For monitoring UDS (Figure 2H), [α-^32^P]dCTP (Revvity; 0.8 µCi/µl) was added. For experiments shown in Figures 1C, 1D and S1E, the “Extract Mixture” was prepared with undepleted NPE, and, where indicated, GG-NER was inhibited by the addition of XPC antibody (1.5 µM final concentration). For all other cell-free repair assays, NPE was first immunodepleted of XPC and the indicated factors (see protocol below), unless indicated otherwise. Compared to IVT assays, the “Extract Mixture” for cell-free repair was further supplemented with a 17.5x stock of “CSB Mixture” and a 11.67x stock of “CSA Mixture” (both used at 1x in the final repair reaction). The “CSB Mixture” contained CSB Buffer (50 mM HEPES pH 7.5, 300 mM NaCl, 10% (v/v) glycerol, and 2 mM DTT) and, where indicated, recombinant hsCSB, xlELOF1, and xlSTK19 variants. The “CSA Mixture” contained CSA Buffer (25 mM HEPES pH 7.5, 150 mM NaCl, 5% (v/v) glycerol, and 2 mM DTT) and, where indicated, recombinant xlCSA-xlDDB1 (or CRL4^CSA^; both can be used interchangeably; not shown) and xlUVSSA. Final repair reactions contained the following concentrations of recombinant TC-NER factors: 200 nM CSB, 100 nM CSA-DDB1, 200 nM ELOF1, 200 nM STK19, and ∼50 nM UVSSA. Instead of using the latter purified protein, *X. laevis* UVSSA variants could be produced in the TnT^®^ SP6 High-Yield Wheat Germ Protein Expression System (Promega) according to manufacturer’s instructions and added directly to the “Extract Mixture”, making up 12% of the final reaction. For the experiment shown in Figure 2F, the cullin inhibitor MLN4924 was supplemented at a final concentration of 200 µM in the “Extract Mixture”.

Once the complete “Extract Mixtures” were prepared, samples were withdrawn for western blot analyses shown in Figures S2C-S2K. Repair reactions were started by combining “Extract Mixture” and “IVT Mixture” such that NPE (supplemented with ATP regenerating system and DTT) represents 50% of the final reaction volume. Reactions were incubated at room temperature for the indicated times, when samples were withdrawn and mixed with seven volumes of IVT Stop Solution containing 0.25 mg/ml RNase A. After 30 min incubation at 37ºC, Proteinase K was added to 1.2 mg/ml, and samples were incubated for 1 h at 37ºC. Plasmids were then purified using AMPure XP Reagent (Beckman), according to manufacturer’s instructions, except that 1.8 volumes of bead solution were added. The DNA was eluted and digested with the appropriate restriction enzyme mix prepared in 1x rCutSmart buffer (NEB) for 1 h at 37ºC. For the UDS assay in Figure 2H, EcoRI-HF and KpnI-HF were added, samples were stopped with 2x IVT Loading Buffer, denatured for 5 min at 95ºC, resolved on a 10% Urea-PAGE gel in 0.8x GTG buffer, vacuum dried, and visualized by autoradiography as described above. For error-free repair experiments, the DNA was digested with XhoI and PmlI and subsequently stopped with Replication Stop Solution (80 mM Tris [pH 8.0], 8 mM EDTA, 0.13% phosphoric acid, 10% Ficoll, 5% SDS, and 0.2% bromophenol blue). DNA products were resolved on a native 0.9% agarose gel in 1x TBE, stained with SYBR Gold (Invitrogen) and imagen on a Typhoon 5 (GE Healthcare). DNA fragments were quantified using ImageJ (NIH) and plotted in GraphPad Prism v.10.2.2. In each independent experiment, the relative error-free repair activity at 120 min shown in Figure 1F was calculated relative to condition I. Relative TC-NER activities in Figure 2A and all subsequent TC-NER assays were determined by calculating the repair activity observed relative to condition I (no TC-NER factor present; given the value 0) and condition II (all wild-type TC-NER factors present; given the value 1), which were included in every independent experiment as reference conditions.

### Antibodies and immunodepletions

All antibodies used in this study are listed in Table S5. Rabbit polyclonal antibodies against the following *X. laevis* peptides were raised and affinity-purified by Biosynth, and used for western blotting and immunodepletions unless otherwise indicated: XPC CT (residues 1049-1062; Ac-CKKGEENHLFPFEKL-OH; used for GG-NER inhibition), XPC NT (amino acids 1-20; H_2_N-MAKRGSSEGAAVAKKKPRKQC-amide; used for depletion and purification), CSB (1357-1370; Ac-CIDGTGVWRLKPEFH-OH; used for western blotting only); CSA (amino acids 380-399; Ac-CHRTHINPAFEDAWSSSEDES-OH),^12^ UVSSA (residues 718-737; Ac-CNRADKSRHEKFANQFNYALN-OH), STK19 (amino acids 1-15; H_2_N-MDRKRKLISDAFKVKC-amide; peptide sequence contains a cysteine 9 to serine substitution), RPB1 (four heptad repeats of the C-terminal domain; Ac-C(YSPTSPS)_4_-amide; used for western blotting), and XPD (residues 741-760; Ac-CLEQLQSEEMLQKIQEIAHQV-OH; used for western blotting). Rabbit polyclonal antibodies against full-length *X. laevis* ELOF1 were prepared by Pocono Rabbit Farm and Laboratory, affinity-purified from serum using the recombinant ELOF1 coupled to AminoLink™ Coupling Resin (Thermo Fisher Scientific) according to manufacturer’s protocol, and used for western blotting and immunodepletions. The following antibodies were used for western blotting: Rabbit polyclonal antibody against *X. laevis* TDG,^37^ and rat monoclonal antibodies targeting RPB1 phospho-Ser5 (clone 3E8) and phospho-Ser2 (clone 3E10), which were a generous gift of Stephen Buratowski.

Depletions were performed with (i) rProtein A Sepharose™ Fast Flow (Cytiva), (ii) Dynabeads™ Protein A for Immunoprecipitation (Thermo), or (iii) Protein A Mag Sepharose Xtra (Cytiva) after equilibration with 1x PBS supplemented with 0.25 mg/ml BSA (and 0.05% Tween in case of (iii)). For (i), five volumes of affinity-purified antibodies (1 mg/ml) were incubated with one volume of beads. For (ii), one volume of affinity-purified antibodies (1 mg/ml) were incubated with two volumes of bead slurry. For (iii), two volumes of affinity-purified antibodies (1 mg/ml) were incubated with one volume of bead slurry. After gentle rotation overnight at 4ºC, beads were washed three times with 1x PBS supplemented with 0.1 mg/ml BSA (and 0.05% Tween in case of (iii)) and three times with egg lysis buffer (ELB; 10 mM HEPES [pH 7.7], 50 mM KCl, 2.5 mM MgCl_2_, and 250 mM sucrose) supplemented with 0.1 mg/ml BSA. Three rounds of depletion were then performed for one hour each at 4ºC by incubating one volume of antibody-bound beads or bead slurry with 5 (i), 0.75 (ii), or 2 (iii) volumes of egg extract. The depleted extracts were collected and immediately used for IVT, GG-NER, or TC-NER assays as described above.

### SDS-PAGE and immunoblotting

Extract samples were stopped with 2x Laemmli sample buffer (120 mM Tris [pH 6.8], 4% SDS, 20% glycerol, 0.02% bromophenol blue, and 10% β-mercaptoethanol), boiled for 2 min at 95ºC, and resolved on 4-15% Mini-PROTEAN® TGX™ Precast Protein Gels or 4-15% Criterion™ TGX™ Precast Midi Protein Gels (Bio-Rad) using Tris-Glycine-SDS Running Buffer (Boston BioProducts). InstantBlue^®^ Protein Stain (Expedeon) was used for coomassie staining. For western blotting, gels were transferred to PVDF membranes (Perkin Elmer). Membranes were blocked with 5% (w/v) non-fat milk in PBST for 30-60 min at room temperature. Primary antibodies were diluted 1:500–1:5,000 in 1x PBST supplemented with 1% BSA and 0.02% sodium azide. After overnight incubation at 4ºC, membranes were extensively washed with 5% (w/v) non-fat milk in PBST at room temperature prior to incubation with horseradish peroxidase (HRP)-conjugated secondary antibodies (Jackson ImmunoResearch) diluted to 1:10,000 in 5% (w/v) non-fat milk in PBST. After secondary antibody incubation for 45-60 min at room temperature, membranes were washed with 1x PBST, incubated with Super-Signal™ West Dura Extended Duration Substrate (Thermo), and imaged on an Amersham™ Imager 600 (GE Healthcare).

### Cloning of TC-NER factors used in cell-free TC-NER

All expression plasmids used in this study are listed in Table S4. Sequences encoding *H. sapiens* STK19 as well as *X. laevis* STK19, UVSSA, CSA, and DDB1 were ordered as codon-optimized gene blocks from Integrated DNA Technologies (IDT). Open reading frames for UVSSA, CSA, and DDB1 were separately cloned into pAceBac1 (pAB1), introducing an N-terminal FLAG tag on UVSSA, leaving CSA untagged, and introducing a His6 tag, followed by a TEV protease cleavage site and an Avi tag on the N-terminus of DDB1. Both CSA and DDB1 plasmids were combined into a single plasmid by I-CeuI restriction digest and Gibson Assembly (NEBuilder^®^ HiFi DNA Assembly Master Mix, New England Biolabs #E2621L). *X. laevis* UVSSA was also cloned into the pF3A backbone (Promega #L5671) for expression in the TnT^®^ SP6 High-Yield Wheat Germ Protein Expression System (Promega). Using Gibson Assembly, open reading frames for both *H. sapiens* and *X. laevis* STK19 were directly cloned into pOPINB (Addgene #41142), introducing a PreScission protease cleavable N-terminal His6 tag, and pOPINK (Addgene #41143), featuring a PreScission protease cleavable N-terminal His6-GST tag. Point mutations and truncations of STK19, UVSSA, and CSA were introduced by “Round-the-Horn” mutagenesis. All plasmids were verified by Sanger or whole plasmid sequencing.

### Protein expression and purification for cell-free transcription and repair

*H. sapiens* CSB variants and *X. laevis* CUL4A-RBX1 were recombinantly expressed in insect cells and purified as described.^12^ *X. laevis* ELOF1 variants were expressed in *E. coli* and purified as described previously.^13^ All purifications were performed at 4ºC unless stated otherwise.

*X. laevis* UVSSA and CSA-DDB1 variants were expressed in insect cells. Bacmids were produced and Sf9 cells (Expression Systems) were maintained and propagated as described previously.^12^ Protein expression was performed in Sf9 (UVSSA) or Hi5 (CSA-DDB1) cells for 72 h at 27ºC (UVSSA) or 20ºC (CSA-DDB1). Harvested cell pellets were resuspended in respective lysis buffer (see below), frozen in liquid nitrogen, and stored at

-80°C prior to purification. Cells were thawed in a water bath, supplemented with one tablet cOmplete™ EDTA-free Protease Inhibitor Cocktail (Roche), lysed by sonication, and cleared by centrifugation for 1 h at 35,000 rpm in a Beckman Ti45 rotor.

The N-terminally FLAG tagged UVSSA was purified using ANTI-FLAG^®^ M2 Affinity Gel (Millipore) equilibrated in UVSSA Lysis Buffer (50 mM HEPES pH 7.5, 500 mM NaCl, 10% glycerol, and 0.1% NP-40). The cleared and filtered (0.8-µm syringe filter) lysate was incubated with the resin for 2 h at 4ºC before washing with UVSSA Wash Buffer (50 mM HEPES pH 7.5, 500 mM NaCl, and 10% glycerol). The resin was equilibrated with UVSSA Buffer (25 mM HEPES pH 7.5, 150 mM NaCl, and 5% glycerol) and eluted with UVSSA Buffer containing 0.2 mg/ml 3x FLAG^®^ Peptide (Sigma). Peak fractions were pooled, DTT was added to a final concentration of 2 mM, the protein was concentrated with a 0.5 ml 10 MWCO spin concentrator (Millipore), frozen in liquid nitrogen, and stored at −80°C.

The CSA-DDB1 heterodimer was purified by applying the cleared and filtered lysate to a 5 ml HisTrap HP column (Cytiva) that was equilibrated in CSA Lysis Buffer (25 mM HEPES pH 7.5, 150 mM NaCl, 10% glycerol, 2 mM 2-mercaptoethanol, and 20 mM imidazole). The column was washed with 4 column volumes (CV) CSA Lysis Buffer, 10 CV CSA High Salt Buffer (CSA Lysis buffer, except 800 mM NaCl), and 4 CV CSA Lysis Buffer. At this point, a 1 ml HiTrap Q HP column (Cytiva) was connected downstream of the HisTrap HP column, and the protein was eluted directly onto the HiTrap Q column with 3 CV CSA Elution Buffer (CSA Lysis buffer, except 300 mM imidazole). The columns were subsequently washed with 3 CV CSA Lysis Buffer prior to the removal of the HisTrap HP column. The protein was eluted from the HiTrap Q column with a linear gradient from CSA Lysis Buffer to CSA High Salt Buffer over 25 CV. Peak fractions were collected, mixed with TEV protease, and dialyzed O/N against CSA Lysis Buffer. The sample was applied to the regenerated and pre-equilibrated HisTrap HP column, and the flow-through was collected, concentrated with a 5 ml 50 MWCO spin concentrator, and loaded onto a Superdex 200 Increase 10/300 GL (Cytiva) equilibrated in CSA Buffer (25 mM HEPES pH 7.5, 150 mM NaCl, 5% (v/v) glycerol, and 2 mM DTT). Peak fractions were pooled, concentrated with 5 ml 50 MWCO spin concentrator (Millipore), frozen in liquid nitrogen, and stored at −80°C.

*H. sapiens* and *X. laevis* STK19 variants as well as GAL4-VP64 and H. sapiens TBP were recombinantly expressed in E. coli OverExpress™ C41(DE3) Chemically Competent Cells (Sigma #CMC0017) or Rosetta™ 2(DE3)pLacI Competent Cells (Millipore #71404) grown at 37ºC in LB media supplemented with the appropriate antibiotics. Typically, 2 l per construct were expressed. At an OD 600 of 0.5-0.7, protein expression was induced with 0.5 mM IPTG, and cultures grown for 18-20 h at 18ºC. Cells were harvested by centrifugation, resuspended in the respective lysis buffer (see below), frozen in liquid nitrogen, and stored at -80°C. To start the purification, cells were thawed in a water bath, supplemented with 1 mg/ml lysozyme and one tablet cOmplete EDTA-free Protease Inhibitor Cocktail (Roche), opened by sonication, and cleared by centrifugation for 1 h at 35,000 rpm in a Ti45 rotor (Beckman).

*X. laevis* STK19 expressed with a PreScission protease cleavable N-terminal His6-GST tag was purified using Glutathione Sepharose 4B resin (Cytiva) equilibrated with STK19 Buffer (25 mM HEPES pH 7.5, 300 mM NaCl, 10% (v/v) glycerol, and 2 mM DTT). After incubation for 1 h at 4ºC, the resin was washed extensively with STK19 Buffer prior to cleavage of the His6-GST tag on the resin with GST-tagged Pre-Scission protease overnight at 4ºC. The released, untagged STK19 was collected and subjected to size-exclusion chromatography on a HiLoad^®^ 16/600 Superdex 75 pg column (Cytiva) equilibrated in STK19 Buffer. Peak fractions were pooled, concentrated, snap-frozen in liquid nitrogen, and stored at -80ºC.

*H. sapiens* and *X. laevis* STK19 variants expressed with a PreScission protease cleavable N-terminal His6 tag were purified using Ni-NTA Superflow resin (Qiagen) equilibrated in STK19 NiNTA Buffer (50 mM HEPES pH 7.5, 300 mM NaCl, 10 mM imidazole, 10% (v/v) glycerol, and 2 mM 2-mercapto-ethanol). Incubation for 1 h at 4ºC was followed by extensive washing of the beads with STK19 NiNTA Wash Buffer (50 mM HEPES pH 7.5, 500 mM NaCl, 20 mM imidazole, 10% (v/v) glycerol, and 2 mM 2-mercaptoethanol). Proteins were eluted with STK19 NiNTA Buffer containing 300 mM imidazole. His-STK19 variants were directly subjected to size-exclusion chromatography on a HiLoad^®^ 16/600 Superdex 75 pg column (Cytiva) equilibrated in STK19 Buffer. Untagged human STK19 was dialyzed against STK19 Buffer after NiNTA elution in the presence of GST-tagged PreScission protease. On the next day, the sample was concentrated and directly subjected to size-exclusion chromatography on a HiLoad^®^ 16/600 Superdex 75 pg column (Cytiva) that had a 1 ml GSTrap™ HP column (Cytiva) attached downstream. Peak fractions were pooled, concentrated with a 5 ml 10 MWCO spin concentrator, frozen in liquid nitrogen, and stored at -80ºC.

The transcription activator GAL4-VP64 and *H. sapiens* TBP were also expressed with an N-terminal His6 tag, but without a PreScission protease site. Purifications were performed analogously and featured the same affinity and size-exclusion steps.

*X. laevis* XPC used in Figure S1 was purified from egg extract (HSS; high-speed supernatant) using a polyclonal antibody that was raised against the XPC N-terminus (see above and Table S5). The antibody was immobilized on rProtein A Sepharose™ Fast Flow (Cytiva) and then incubated with HSS overnight at 4°C. The beads were washed with Wash Buffer (25 mM HEPES pH 7.5, 0.1 mM EDTA, 12.5 mM MgCl_2_, 10% glycerol, 0.1 mM DTT and 0.01% NP-40) supplemented with 400 mM KCl followed by Wash Buffer supplemented with 200 mM KCl. The endogenous XPC protein was eluted with Wash Buffer containing 200 mM KCl and 1.5 mg/ml XPC NT peptide (Biosynth; H_2_N-MAKRGSSEGAAVAKKKPRKQC-amide). The eluted sample was subjected to size exclusion chromatography on a Superose 6 Increase column (Cytiva) equilibrated with XPC SEC Buffer (25 mM HEPES pH 7.5, 150 mM NaCl, 10% glycerol, and 2 mM DTT). Peak fractions were pooled, concentrated with a 5 ml 10 MWCO spin concentrator (Millipore), frozen in liquid nitrogen, and stored at −80°C.

### AF-M Model Generation

Unless otherwise stated, AlphaFold multimer (AF-M)^38^ structure predictions were generated using ColabFold 1.5^39^ running AF-M with v2.3 weights, 1 ensemble, 3 recycles, templates enabled, without dropout, and with maximum Multiple Sequence Alignments MSA depth settings (max_seq = 508, max_extra_seq = 2048). MSAs (paired and unpaired) were provided to AlphaFold multimer via the MMSeq2^40^ API built into ColabFold.

### AF-M Model Analysis

Analysis of structural predictions generated by AF-M was performed using python scripts as previously described.^27,41^ Briefly, confident interchain residue contacts were extracted from structures by identifying proximal residues (distance <5 Å) where both residues have pLDDT values >50 and PAE score <15 Å. All subsequent downstream analysis of interface statistics (average pLDDT, average PAE) were calculated using data associated with these inter-residue pairs (contacts). Average interface pLDDT values above 70 are generally considered confident.^42^ The average models score was calculated by averaging how many independent AF-M models predicted a contact across all unique inter-residue contact pairs. This number was then normalized by dividing the number of models run to produce a final score that ranges from 0 (worst) to 1 (best). An average models score above 0.5 is considered confident. pDOCKQ estimates of interface accuracy scores were calculated independently of the contact analysis described above using code from ^43^. pDOCKQ values above 0.23 are considered confident.

### SPOC Analysis

A random forest classifier (Structure Prediction and Omics Classifier) SPOC trained to distinguish biologically relevant interacting protein pairs from non-relevant interaction pairs was run on AF-M predictions as described previously.^27^ For every interaction evaluated, it generates a score that can range from 0 (worst) to 1 (best). Higher scores indicate that AF-M interface metrics and several types of externally sourced biological data are consistent with the existence of the binary interaction tested. SPOC scores above 0.5 are generally associated with high confidence interaction predictions.

### Biolayer interferometry (BLI)

DNA binding experiments were performed on an Octet RED384 system (Sartorius). A 5’-biotinylated 14mer (5’–Bio–TAT GGA CAG CAA GC–3’) was mixed and annealed with a complementary 14mer (5’–GCT TGC TGT CCA TA–3’) in Annealing Buffer (10 mM Tris pH 7.5, 50 mM NaCl, and 1 mM EDTA). The DNA duplex was captured at a concentration of 1.25 µg/ml in BLI Buffer (ELB containing 0.05% Tween-20 and 5 mg/ml BSA) on an Octet^®^ Streptavidin (SA) Biosensor (Sartorius). His-tagged *X. laevis* STK19 variants were diluted to 500 nM in BLI Buffer. Binding to the immobilized DNA duplex was conducted in parallel in Octet^®^ 384 Well Tilted Bottom Plates (Sartorius) at 30ºC, shaking at 1,000 rpm, using the following steps: 120 s baseline (BLI Buffer), 120 s sample (DNA duplex) loading, 120 s baseline (BLI Buffer), 300 s analyte (STK19) association, 600 s dissociation. The data was processed in Octet Analysis Studio 13.0 and analyzed with Prism 10 (GraphPad).

### *In vitro* ubiquitination assays

*X. laevis* CRL4^CSA^ complexes were assembled by incubating equimolar amounts of purified CSA-DDB1 variants with CUL4A-RBX1 for 15 min at room temperature. Complexes were then neddylated using the NEDD8 Conjugation Initiation Kit (R&D Systems) according to the manufacturer’s instructions, except using reduced final concentrations for Uba3 (0.5x), UbcH12 (0.5x), and NEDD8 (0.33x). After 30 min incubation at room temperature, ubiquitination reactions were prepared in Ubiquitination Buffer (40 mM Tris pH 7.5, 10 mM MgCl_2_, 0.6 mM DTT), and contained the following proteins: 100 nM E1 enzyme (R&D Systems), 1 µM E2 enzyme (UBE2E1 for UVSSA monoubiquitination or UBE2D2 for CSB polyubiquitination; UBPBio), 1 µM of the indicated neddylated CRL4^CSA^ variant, 25 µM ubiquitin, and 400 nM substrate (UVSSA or CSB). Reactions were started by the addition of 10 mM ATP, incubated at room temperature for the indicated times, stopped with 2x Laemmli sample buffer, and analyzed by SDS-PAGE and western blotting. Samples of the UVSSA monoubiquitination assay were resolved on a 7.5% gel, whereas CSB polyubiquitination was analyzed on a 4-15% SDS-PAGE gel, as described above.

### Cloning of human TC-NER factors

All expression plasmids used in this study are listed in Table S4.Sequences encoding *H. sapiens* TC-NER factors were cloned into the 438-series vectors or the 1-series vectors (MacroLab vectors, UC Berkeley). Initial ORFs were obtained by PCR amplification from cDNA or by synthesis of gBlocks (IDT DNA Technologies). The ORFs were subsequently introduced in the respective vectors using ligation-independent cloning. CSB was tagged with an N-terminal His6 tag, followed by a TEV protease cleavage site (vector 438-B). CSB, amplified from cDNA, had three amino acid variants compared to the canonical isoform (V1097M, G1213R, and R1413Q). None of the three amino acid variants are resolved in our structure or impact reconstitution of TC-NER. UVSSA was tagged with an N-terminal His6 tag, followed by a TEV protease cleavage site (vector 438-B). CSA was cloned with no tag (vector 438-A). DDB1 was tagged with an N-terminal His6 tag, followed by a TEV protease cleavage site (vector 438-B). The DDB1 and CSA vectors were subsequently combined into a multi-ORF plasmid using ligation-independent cloning. DDA1 was tagged with an N-terminal His6 tag, maltose binding protein (MBP) tag, a 10-residue asparagine linker, and a TEV protease cleavage site.

### Protein expression and purification of *S. scrofa* RNA polymerase II and *H. sapiens* TC-NER factors

Plasmid propagation and bacmid isolation were performed in Escherichia coli DH5a (plasmid propagation) or Escherichia coli DH10 EMBacY (bacmid isolation). Recombinant protein expression in insect cells was performed using Sf9, Sf21, and Hi5 cell lines as described.^44^ Recombinant expressions in insect cells were harvested after 72 h. Recombinant protein expressions in insect cells or *E. coli* were centrifuged to harvest. Cell pellets were resuspended in respective lysis buffer (see below), flash-frozen, and stored at -80°C prior to purification.

*H. sapiens* CSB, UVSSA, and CSA-DDB1 were recombinantly expressed in insect cells. DDA1 and ELOF1 were recombinantly expressed in *E. coli* BL21 (DE3) RIL. Specifically, 4 L or 6 L of 2X YT media were inoculated with DDA1 or ELOF1 overnight culture under appropriate antibiotic selection. At OD 600, protein expression was induced with 1 mM of IPTG. Cells were grown at 37°C for three hours before harvest via centrifugation.

*Sus scrofa* RNA polymerase II was purified from pig thymus as described.^45^ *H. sapiens* ELOF1 was purified essentially as described.^15^ The expression and purification of *H. sapiens* STK19 was described above.

All *H. sapiens* TC-NER factor protein purifications were performed at 4°C, unless otherwise specified. The purity of protein preparations was assessed using SDS-PAGE with NuPAGE 4-12% Bis-Tris protein gels (Invitrogen) followed by OneStep Blue (Biotium) staining. Cells were thawed in a water bath. Following cell lysis by sonication, the lysate was clarified using centrifugation and subsequent ultracentrifugation. The supernatant was then filtered using 0.8-µm syringe filters. All protein concentrations were determined by measuring absorption at 280 nm and using the predicted extinction coefficient for the protein(s).

For *H. sapiens* CSB, the filtered supernatant was applied to a 5 mL HisTrap HP column (Cytiva Life Sciences) equilibrated in lysis buffer (500 mM NaCl, 20 mM Na-HEPES pH 7.4, 10% (v/v) glycerol, 30 mM imidazole pH 8.0, and 5 mM 2-mercaptoethanol, 0.284 µg ml−1 leupeptin, 1.37 µg ml−1 pepstatin A, 0.17 mg ml−1 PMSF, and 0.33 mg ml−1 benzamidine). The column was washed with 5 column volumes (CV) of lysis buffer, followed by 20 CV of High Salt buffer (1 M NaCl, 20 mM Na-HEPES pH 7.4, 10% (v/v) glycerol, 30 mM imidazole pH 8.0, and 5 mM 2-mercaptoethanol, 0.284 µg ml−1 leupeptin, 1.37 µg ml−1 pepstatin A, 0.17 mg ml−1 PMSF, and 0.33 mg ml−1 benzamidine). The column was subsequently washed with 5 CV of lysis buffer and the protein was eluted with a 20 CV linear gradient from 0% to 100% Elution buffer (lysis buffer with 500 mM imidazole). Peak fractions were pooled, mixed with TEV protease and dialyzed overnight in 7 kDa MWCO SnakeSkin dialysis tubing (Thermo Scientific) against dialysis buffer (500 M NaCl, 20 mM Na-HEPES pH 7.4, 10% (v/v) glycerol, 50 mM imidazole pH 8.0, and 5 mM 2-mercaptoethanol, 0.284 µg ml−1 leupeptin, 1.37 µg ml−1 pepstatin A, 0.17 mg ml−1 PMSF, and 0.33 mg ml−1 benzamidine). The protein was then applied to tandem HisTrap/ HiTrap Heparin HP (5 ml each) columns (Cytiva) equilibrated in dialysis buffer. The columns were washed with 5 CV of dialysis buffer after which the HisTrap column was removed. The protein was eluted off the HiTrap Heparin HP column using a 20 CV long linear gradient from 0% dialysis buffer to 100% high salt buffer. Peak fractions were pooled and concentrated with 100 kDa MWCO Amicon Ultra Centrifugal Filters (Merck) and applied to a Superdex 200 Increase 10/300 GL (Cytiva) column equilibrated in gel filtration buffer (500 mM NaCl, 20 mM Na-HEPES pH 7.4, 10% (v/v) glycerol, and 1 mM TCEP). Fractions containing CSB were concentrated with a 100 kDa MWCO Amicon Ultra Centrifugal Filters. Protein concentration was determined as described and CSB was aliquoted, flash-frozen, and stored at −80°C.

For *H. sapiens* UVSSA, the filtered supernatant was applied to a 5 ml HisTrap HP column equilibrated in lysis buffer. The column was washed with 5 CV of lysis buffer (500 mM NaCl, 20 mM Na-HEPES pH 7.4, 10% (v/v) glycerol, 30 mM imidazole pH 8.0, and 5 mM 2-mercaptoethanol, 0.284 µg ml−1 leupeptin, 1.37 µg ml−1 pepstatin A, 0.17 mg ml−1 PMSF, and 0.33 mg ml−1 benzamidine), followed by 20 CV of high salt buffer (1 M NaCl, 20 mM Na-HEPES pH 7.4, 10% (v/v) glycerol, 30 mM imidazole pH 8.0, and 5 mM 2-mercaptoethanol, 0.284 µg ml−1 leupeptin, 1.37 µg ml−1 pepstatin A, 0.17 mg ml−1 PMSF, and 0.33 mg ml−1 benzamidine). The column was washed with 5 CV of Lysis buffer and 5 CV of low salt buffer (150 mM NaCl, 20 mM Na-HEPES pH 7.4, 10% (v/v) glycerol, 30 mM imidazole pH 8.0, and 5 mM 2-mercaptoethanol, 0.284 µg ml−1 leupeptin, 1.37 µg ml−1 pepstatin A, 0.17 mg ml−1 PMSF, and 0.33 mg ml−1 benzamidine). After washing, a HiTrap Q HP (5 ml) column (Cytiva, equilibrated in low salt buffer) was attached to the HisTrap HP column. The protein was eluted onto the HiTrap Q HP with a 10 CV step gradient of 100% elution buffer (lysis buffer supplemented with 500 mM imidazole). The HisTrap HP column was then removed, and the HiTrap Q column was washed with 5 CV of low salt buffer. The protein was eluted off the HiTrap Q HP column using a 20 CV long linear gradient from 0% low salt buffer to 100% high salt buffer. Peak fractions were pooled, mixed with TEV protease, and dialyzed overnight in 7 kDa MWCO SnakeSkin dialysis tubing (Thermo Scientific) against low salt buffer. The protein was then applied to a HiTrap S HP (5 ml) column (GE Healthcare) equilibrated in low salt buffer. The column was washed with 10 CV of low salt buffer and the protein was eluted off the HiTrap S HP column using a 20 CV linear gradient from 0% low salt buffer to 100% high salt buffer. Peak fractions were pooled and concentrated with 50 kDa MWCO Amicon Ultra Centrifugal Filters (Merck) and applied to a Superdex 200 Increase 10/300 GL (Cytiva) column equilibrated in 500 mM NaCl, 20 mM Na-HEPES pH 7.4, 10% (v/v) glycerol, and 1 mM TCEP. Fractions containing UVSSA were concentrated with 50 kDa MWCO Amicon Ultra Centrifugal Filters (Merck). Protein concentration was determined as described and UVSSA was aliquoted, flash-frozen, and stored at −80°C.

For *H. sapiens* CSA-DDB1, the filtered supernatant was applied to a 5-ml HisTrap HP column (GE Healthcare) equilibrated in lysis buffer. The column was washed with 5 CV of Lysis buffer (400 mM NaCl, 20 mM Na-HEPES pH 7.4, 10% (v/v) glycerol, 30 mM imidazole pH 8.0, and 5 mM 2-mercaptoethanol, 0.284 µg ml−1 leupeptin, 1.37 µg ml−1 pepstatin A, 0.17 mg ml−1 PMSF, and 0.33 mg ml−1 benzamidine), followed by 20 CV of High Salt buffer (1 M NaCl, 20 mM Na-HEPES pH 7.4, 10% (v/v) glycerol, 30 mM imidazole pH 8.0, and 5 mM 2-mercaptoethanol, 0.284 µg ml−1 leupeptin, 1.37 µg ml−1 pepstatin A, 0.17 mg ml−1 PMSF, and 0.33 mg ml−1 benzamidine). The column was washed with 5 CV of Lysis buffer and 5 CV of low salt buffer (150 mM NaCl, 20 mM Na-HEPES pH 7.4, 10% (v/v) glycerol, 30 mM imidazole pH 8.0, and 5 mM 2-mercapto-ethanol, 0.284 µg ml−1 leupeptin, 1.37 µg ml−1 pepstatin A, 0.17 mg ml−1 PMSF, and 0.33 mg ml−1 benzamidine). After washing, a HiTrap Q HP (5 ml) column (GE Healthcare) equilibrated in low salt buffer was attached to the 5-ml HisTrap HP column. The protein was eluted onto the HiTrap Q HP with a 10 CV long step gradient of 100% Elution buffer (Lysis buffer supplemented with 500 mM imidazole). The HisTrap HP column was then removed and the HiTrap Q column was then washed with 5 CV of Low salt buffer. The protein was eluted off the HiTrap Q HP column using a 20 CV long linear gradient from 0% Low salt buffer to 100% High salt buffer. Peak fractions were pooled, mixed with TEV protease and dialysed overnight in 7 kDa MWCO SnakeSkin dialysis tubing (Thermo Scientific) against Dialysis buffer (400 mM NaCl, 20 mM Na-HEPES pH 7.4, 10% (v/v) glycerol, 50 mM imidazole pH 8.0, and 5 mM 2-mercaptoethanol, 0.284 µg ml−1 leupeptin, 1.37 µg ml−1 pepstatin A, 0.17 mg ml−1 PMSF, and 0.33 mg ml−1 benzamidine). The flowthrough containing the complex was pooled and concentrated with 100 kDa MWCO Amicon Ultra Centrifugal Filters (Merck) and applied to a Superdex 200 Increase 10/300 GL (Cytiva) column equilibrated in 400 mM NaCl, 20 mM Na-HEPES pH 7.4, 10% (v/v) glycerol, and 1 mM TCEP. Fractions containing the complex were concentrated with 100 kDa MWCO Amicon Ultra Centrifugal Filters (Merck). Protein concentration was determined as described and CSA-DDB1 was aliquoted, flashfrozen, and stored at −80°C.

For *H. sapiens* DDA1, the filtered supernatant was applied to a 5-ml HisTrap HP column (GE Healthcare) equilibrated in lysis buffer (500 mM NaCl, 20 mM Na-HEPES pH 7.4, 10% (v/v) glycerol, 30 mM imidazole pH 8.0, and 5 mM 2-mercapto-ethanol, 0.284 µg ml−1 leupeptin, 1.37 µg ml−1 pepstatin A, 0.17 mg ml−1 PMSF, and 0.33 mg ml−1 benzamidine). The column was washed with 5 CV of lysis buffer followed by 20 CV of high salt buffer, and 5 CV of lysis buffer. A self-packed XK column (Cytiva) with 15 mL of amylose resin (New England Biolabs) was attached to the HisTrap column which was then equilibrated into Lysis buffer. Sample was directly eluted onto the Amylose column with a 10 CV long step gradient of 100% elution buffer (lysis buffer supplemented with 500 mM imidazole). The HisTrap HP column was removed, and the amylose column was then washed with 5 CV lysis buffer. The protein was eluted off the amylose column using an amylose elution buffer (lysis buffer supplemented with 116.9 mM maltose). Peak fractions were pooled, mixed with TEV protease and dialysed overnight in 7 kDa MWCO SnakeSkin dialysis tubing (Thermo Scientific) against lysis buffer (500 mM NaCl, 20 mM Na-HEPES pH 7.4, 10% (v/v) glycerol, 50 mM imidazole pH 8.0, and 5 mM 2-mercaptoethanol, 0.284 µg ml−1 leupeptin, 1.37 µg ml−1 pepstatin A, 0.17 mg ml−1 PMSF, and 0.33 mg ml−1 benzamidine). The dialyzed sample was subsequently applied to a 5 mL HisTrap column equilibrated in lysis buffer. The flow-through containing DDA1 was pooled and concentrated with 10 kDa MWCO Amicon Ultra Centrifugal Filters (Merck) and applied to a HiLoad 16/600 Superdex 75 (Cytiva) column equilibrated in 500 mM NaCl, 20 mM Na-HEPES pH 7.4, 10% (v/v) glycerol, and 1 mM TCEP. Fractions containing DDA1 were concentrated with 10 kDa MWCO Amicon Ultra Centrifugal Filters (Merck). Protein concentration was determined as described and DDA1 was aliquoted, flash-frozen, and stored at −80°C.

### Complex preparation for cryo-EM

All concentrations refer to the final concentrations in the transcription reaction. RNA (5’–AUCGAGAGGA–3’) and template DNA (5’–GAG GTC ACT CCA GTG AAT TCG AGC TCG CAA CAA TGA GCA CAT TCG CTC TGC TCC TTC TCC CAT CCT CTC GAT GGC TAT GAG ATC AAC TAG–3’) were mixed and annealed as described.^11^ 80 pmol nM of DNA·RNA hybrid were incubated with 80 pmol *S. scrofa* RNA polymerase II and incubated for 10 min on ice. 1040 pmol non-template DNA (5’– CTA GTT GAT CTC ATA TTT CAT TCC TAC TCA GGA GAA GGA GCA GAG CGA ATG TGC TCA TTG TTG CGA GCT CGA ATT CAC TGG AGT GAC CTC–3’) was added to the mixture and again incubated for 10 min on ice. A factor mix was prepared separately with 2400 pmol each of UVSSA, CSB, and CSA-DDB1, as well as 3200 pmol each of STK19, ELOF1, and DDA1. The factor mix was incubated for 10 min on ice. The factor mix was subsequently added to the elongation complex and incubated for 10 min on ice. ADP·BeF_3_, water and compensation buffer were added to adjust to a final concentration of buffer components of 330 mM NaCl, 20 mM Na·HEPES pH 7.4, 5 % (v/v) glycerol, 0.3 mM ADP·BeF_3_, and 1 mM TCEP. The reaction was incubated for 10 min on ice. The sample was dialyzed for 3 hours into a final buffer of 100 mM NaCl, 20 mM HEPES, pH 7.4, 3 mM MgCl_2_, 5% (v/v) glycerol, and 1 mM TCEP. The sample was centrifuged for 10 min at 21,300*g* and applied to a Superose 6 Increase 3.2/300 (Cytiva) on an Äkta pure 25 with Micro kit (Cytiva). 50 μL fractions were collected. Peak fraction samples were applied to NuPAGE 4-12% Bis-Tris gels (Invitrogen) and run in 1X MES buffer for 30 min at 200 V to assess complex formation. The gel was stained with One-Step Blue Protein Gel Stain (Biotum) and imaged. Relevant peak fractions were individually crosslinked with 0.1% (v/v) glutaraldehyde for 10 min on ice and then quenched with 8 mM aspartate and 2 mM lysine for 10 min on ice. The reactions were transferred to a Slide-A-Lyzer Mini Dialysis Unit 20 K MWCO (Thermo) and dialyzed against buffer containing 100 mM NaCl, 20 mM Na-HEPES pH 7.4, and 1 mM TCEP for 3 h at 4°C.

Complex concentrations were quantified by absorbance at 280 nm. The molar extinction coefficient of the complex was obtained by summing the molar extinction coefficients of all individual components. The fraction with the nominal highest concentration (∼600 nM) was selected for analysis by cryo-EM. Quantifoil UltrAufoil R 2/2, 200 Mesh, Au grids were glow discharged for 120 s at 30 mA and 0.38 mBar with 10 s hold time at 0.38 mBar using a Pelco Easiglow plasma discharge system. 2 μL of sample were applied on each side of the grid, incubated for 10 s, blotted with Ted Pella standard Vitrobot filter paper for 3 s with blot force 8 and vitrified by plunging into liquid ethane using a Vitrobot Mark IV (FEI Company), operated at 4°C and 100% humidity. Sample application from both sites and sample incubation for 8 s has consistently resulted in better ice quality for transcription elongation complexes compared to single-sided sample application.

### Cryo-electron microscopy and image processing

Cryo-EM data were collected on a Thermo Fisher Scientific Titan Krios operated at 300 keV equipped with a Falcon 4 and a Selectris energy filter. Data acquisition was automated using EPU at a nominal magnification of 130,000x, corresponding to a pixel size of 0.94 Å in nanoprobe EFTEM mode. Movies consisting of 63 frames were collected in counted mode with an exposure time of 5.53 s. The electron flux was 9.48 e^−^ Å^-2^ s^-1^ with a total dose of 52.4 e^−^ Å^-2^. Image processing and analysis were performed with cryoSPARC (v 4.4.1) using default parameters, unless stated otherwise.

Movies were aligned using patch motion correction followed by contrast transfer function (CTF) estimation in cryoSPARC. Particles were picked by blob-based automatic picking, resulting in 3,742,711 particles from 15,521 micrographs. Particles were extracted with a Fourier binned box size of 300 pixels (pixel size of 1.47 Å). All classifications and refinements were conducted in cryoSPARC. Volumes employed for masking of areas of interest were generated by low-pass filtering the regions of interest to 10-15 Å and then using cryoSPARC to expand the volume containing the area of interest by 1-3 hard pixels and 3-7 soft pixels.

Three *ab initio* models were generated from a subset of 20,000 particles. The *ab initio* models were then used to select for particles that contain Pol II and TC-NER factors via heterogeneous refinements and 3D classification, resulting in a subset of about 1,391,445 particles. The selected particles were subsequently 3D classified to remove low-resolution particles. A non-uniform refinement was performed, and the particles (1,123,600 particles) were recentered and re-extracted (box size of 468 pixels with a pixel size of 0.94 Å). The re-extracted particles were refined. Local and global CTF refinement and reference-based motion correction was completed. The subsequent refined particles were further classified using 3D classifycation to improve occupancy of DDB1 and Pol II subunits. Particle duplicates were removed, and the particles were again refined. Subsequent local CTF refinement and non-homogeneous refinement resulted in the 1.9 Å reconstruction of the TC-NER complex with STK19 and DDA1 (map i), close to the nominal Nyquist frequency of 1.88 Å. Refinements of diverse areas of the map were performed including local refinements of the TC-NER factors (map ii), CSB (map iii), STK19 (map iv), DDB1-DDA1 (map v), and UVSSA-DDB1-DDA1 (map vi), RPB4/7 (map vii), and a 3D classification and subsequent local refinement for DDA1 (map viii). With help of an initial model of the complex, a composite map (map ix) of map i, iii, iv, vii, and viii was generated using PHENIX (v 1.20.1).

### Model building and refinement

Structures of the TC-NER complex with ELOF1 (PDB ID 8B3D), were rigid body docked into the overall map. AlphaFold-Multimer structures corresponding to CSA-CSB, DDB1-CSA-UVSSA, CSB, STK19, DDA1-DDB1, and STK19-RPB1 predictions were superposed onto the initial TC-NER structure and used to further built/replace chains of the initial model to complete the structure of the TC-NER complex with STK19 and DDA1. Density in the active site of Pol II allowed unambiguous assignment of the DNA register by defining purine and pyrimidine bases in the DNA·RNA hybrid. Identification of the register was conducted in map i. We additionally observed two additional densities next to the metal A and docked water molecules into these densities. We note additional density for three amino acids bound to ATPase lobe 1 of CSB that we could not unambiguously assign. The atomic model was locally adjusted and refined using ISOLDE (v1.7.1) with help of all local refinements. The overall atomic model was subsequently real space refined in PHENIX against map ix. Refinement statistics are reported in Table S2. Additional information on input structural models and model confidence is given in Table S3.

### Figure generation

All structure figures were generated in UCSF ChimeraX. Angular distribution plots were generated using the available Warp tool. FSC curves were generated in cryoSPARC and adjusted in Adobe Illustrator.

**Figure S1.**
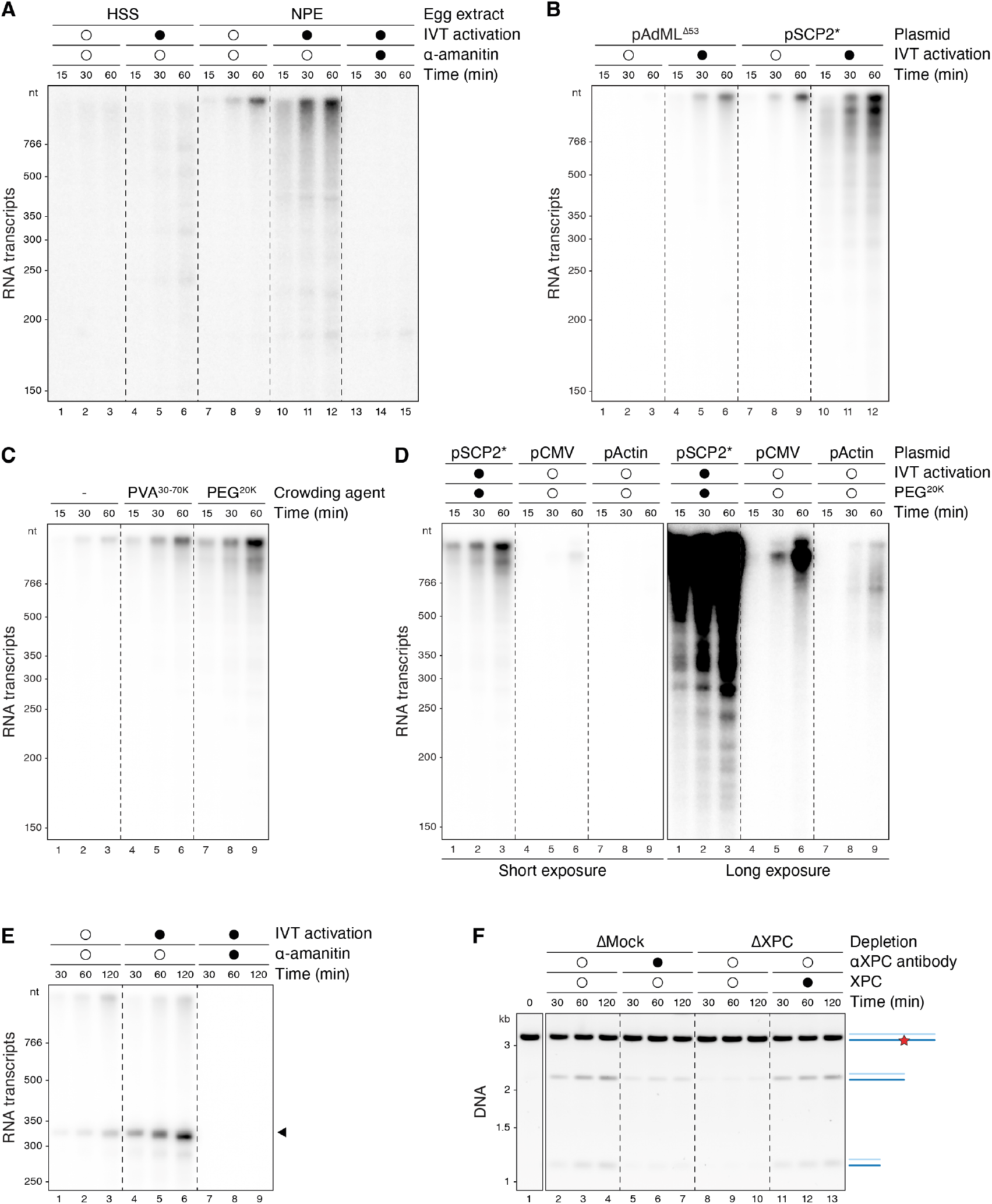
Optimization of transcription efficiency in frog egg extracts. **(A)** Plasmid pAdML^Δ53^ (adenovirus major late promoter flanked by UAS sites) was incubated with total egg lysate (HSS, high speed supernatant) or NPE containing [α-^32^P]UTP that were also optionally supplemented with the transcription activator GAL4-VP64 and TBP (for *in vitro* transcription (IVT) activation) and 2 µM α-amanitin. At the indicated times, RNA was recovered, separated on a Urea-PAGE gel, and subjected to autoradiography. **(B)** Cell-free transcription was carried out in NPE as in (A), comparing plasmids containing the adenovirus major late promoter and a modified super core promoter 2, SCP2* (see Methods). **(C)** Cell-free transcription was carried out in NPE as in (A), with the indicated crowding agents being added to a final concentration of 1% (v/v). **(D)** Comparison of our optimal inducible transcription condition (pSCP2* substrate, IVT activation, and 1% PEG^20K^) to transcription from the CMV and endogenous actin promoters. Plasmids pCMV and pActin were added to NPE without any supplements as in ^25^. Short and long exposures of the autoradiograph are shown. **(E)** The reactions described in Figure 1C were supplemented with [α-^32^P]UTP and used to monitor transcription, demonstrating that IVT activation strongly stimulated transcription, and that α-amanitin inhibited transcription in this experiment. **(F)** NPE supports GG-NER. Plasmid containing a cisplatin 1,3-GTG intrastrand crosslink was incubated in mock-depleted NPE supplemented with buffer or inhibitory XPC antibody, or in NPE depleted of XPC that was optionally supplemented with XPC protein purified from egg extract. DNA was recovered, and PmlI site regeneration was monitored (as depicted in Figure 1A). DNA was separated by agarose gel electrophoresis and visualized using SYBR Gold, which showed that XPC antibody or XPC depletion inhibited error-free repair, with the latter defect being reversed with purified XPC protein.

**Figure S2.**
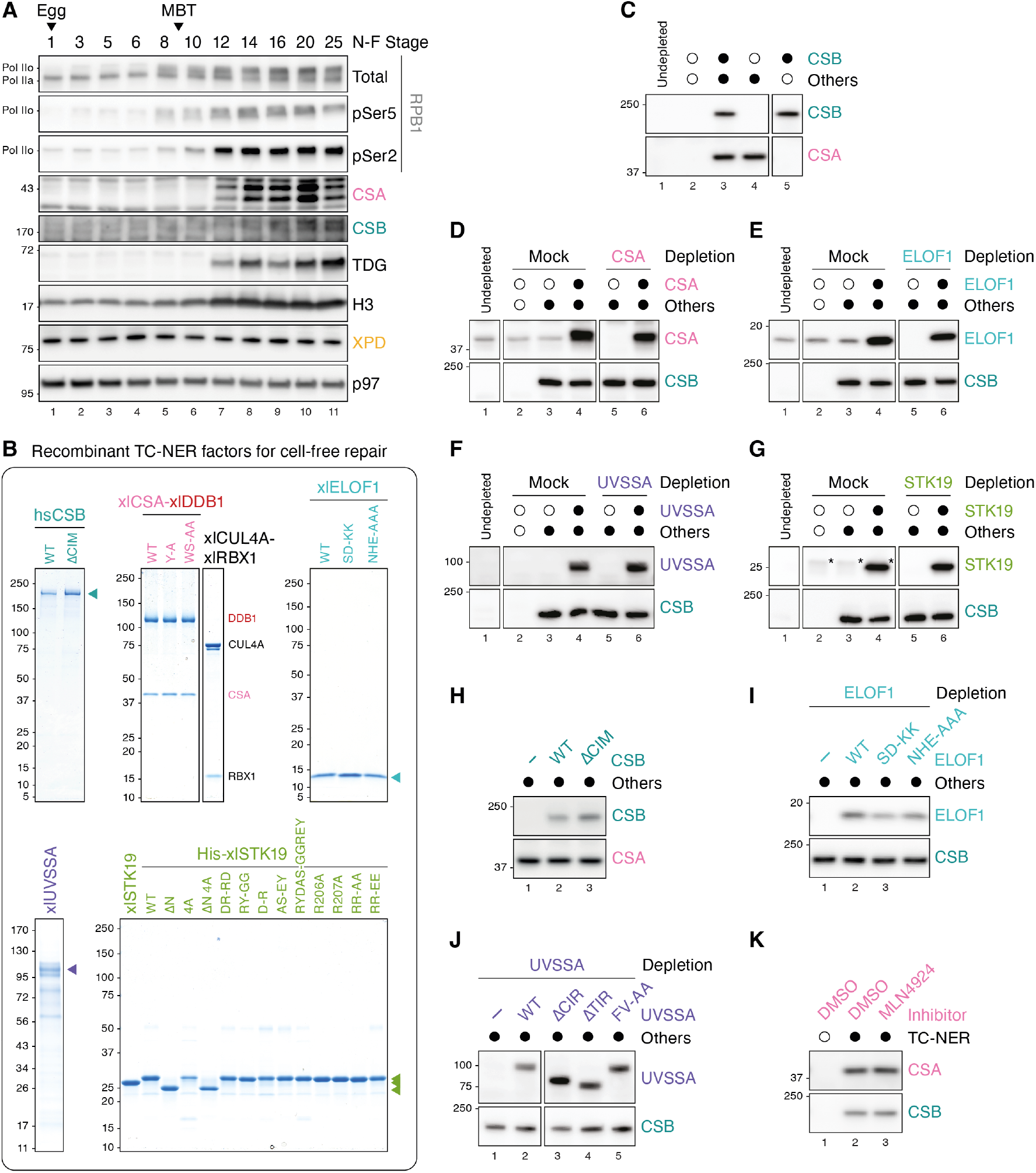
Characterization and purification of TC-NER factors. **(A)** Frog eggs were fertilized *in vitro*, and at the indicated stages, embryo extracts^37^ were blotted for the indicated proteins. Transcription activation during the mid-blastula transition (MBT) was detected by hyperphosphorylation of the RPB1 subunit of Pol II. Protein levels of the TC-NER factors CSA and CSB followed a similar trend as the developmentally regulated TDG (Thymine DNA Glycosylase),^37^ being only detectable after the MBT. In contrast, p97 and the TFIIH subunit XPD were similarly abundant at all stages .**(B)**Proteins used for cell-free TC-NER assays throughout the paper were separated by SDS-PAGE and stained with Coomassie blue. *H. sapiens* CSB was used throughout our study because it activated TC-NER as efficiently as *X. laevis* CSB (not shown) while exhibiting slightly better protein stability. For all other TC-NER factors, *X. laevis* proteins were used. **(C)** Representative western blot of the extracts used for conditions III-IV in Figure 2A. Error-free repair data from extracts in lanes 4 (condition III) and 5 (condition IV) were quantified relative to repair in lanes 2 and 3. The CSA blot is representative of the “Other” TC-NER factors, other than CSB. Note that CSB is absent in NPE and therefore not detected in lane 1. **(D-G)** Representative western blots of the extracts used for conditions V-VIII in Figure 2A and conditions III-X in Figure 2B. In each panel, repair data from extracts in lane 3 was quantified relative to repair in extracts in lanes 2 and 4, producing data points for conditions V-VIII in Figure 2A. Lanes 5 and 6 correspond to the depletion and add-back samples for CSA, ELOF1, UVSSA, and STK19, respectively, shown in conditions III-X in Figure 2B. In addition to blotting for CSA (D), ELOF1 (E), UVSSA (F), and STK19 (G), we also blotted for CSB to represent all the “Other” TC-NER factors in each reaction. Asterisk (*) in G indicates a non-specific band. Note that endogenous levels of UVSSA (F, lane 1) and STK19 (G, lane 1) are below the detection limit of our antibodies. **(H-K)** Representative western blots of the extracts used in experiments shown in Figures 2C-2F.

**Figure S3.**
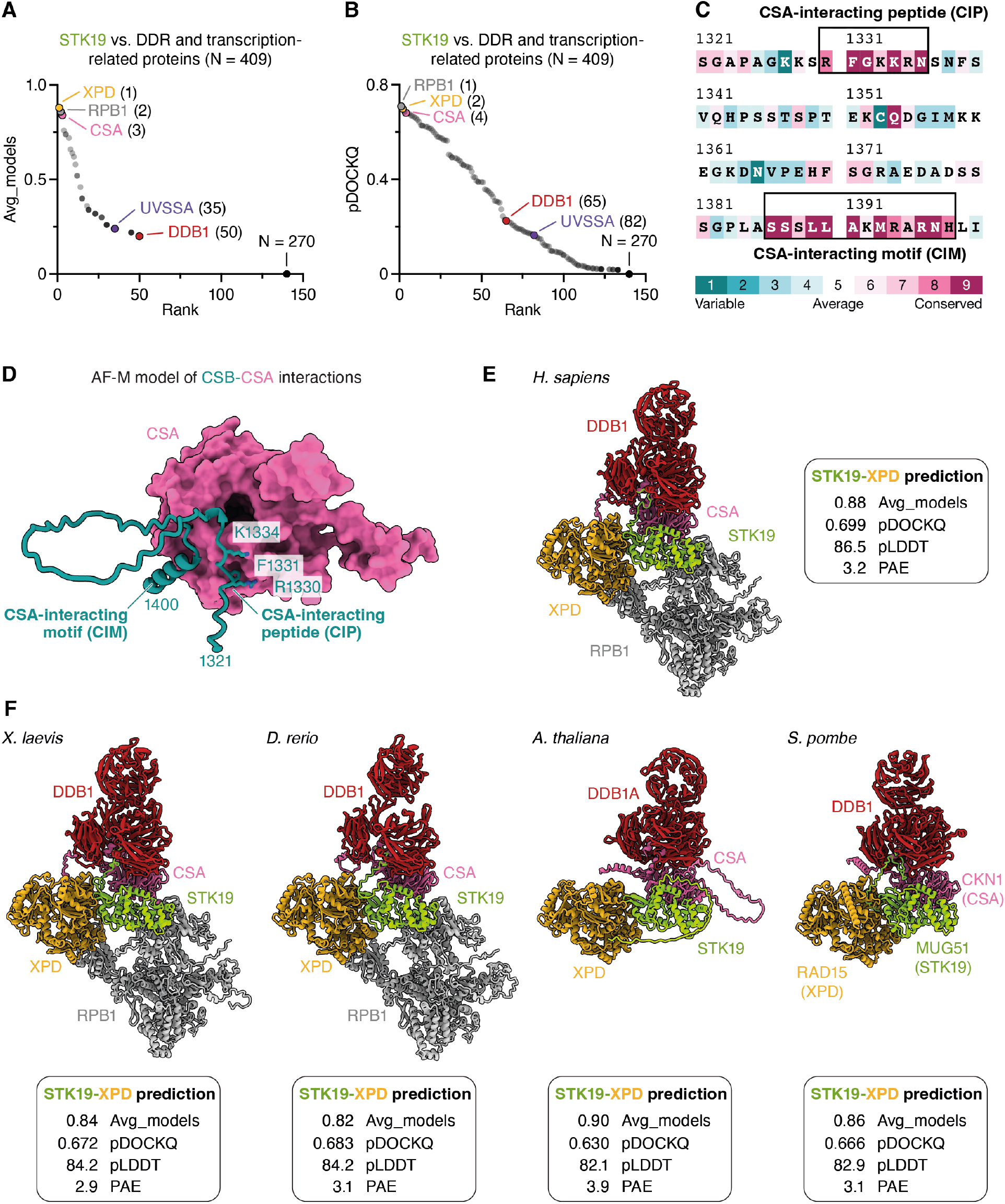
Structure prediction results. **(A-B)** The ∼400 binary structure predictions shown in Figure 3A were ranked based on avg_models (A), a confidence metric that quantifies the agreement among the five different AF-M models, or pDOCKQ (B). **(C)** Sequence conservation of human CSB residues 1321-1400 calculated in ConSurf.^46^ The CSA-interacting peptide (CIP) and the CSA-interacting motif (CIM) of CSB are highly conserved, whereas residues flanking these regions are more variable. **(D)** AF-M prediction for the interaction between CSB and CSA within the folded TC-NER complex (Figures 3B and 3C). CSA is shown in surface representation, and CSB residues 1321-1400 are depicted in cartoon representation. Key residues of CSB in the CSA-interacting peptide (CIP) identified here are shown as side chains. **(E-F)** Composite structure prediction of STK19, DDB1, CSA, XPD, and RPB1 complex in human (E) and other organisms (F). STK19 was folded separately with CSA-DDB1, RPB1, or XPD, and the resulting models were aligned on STK19 to show a composite complex. No interaction was predicted for STK19 with RPB1 in *A. thaliana* and *S. pombe*. The confidence metrics shown refer to the binary STK19-XPD prediction.

**Figure S4.**
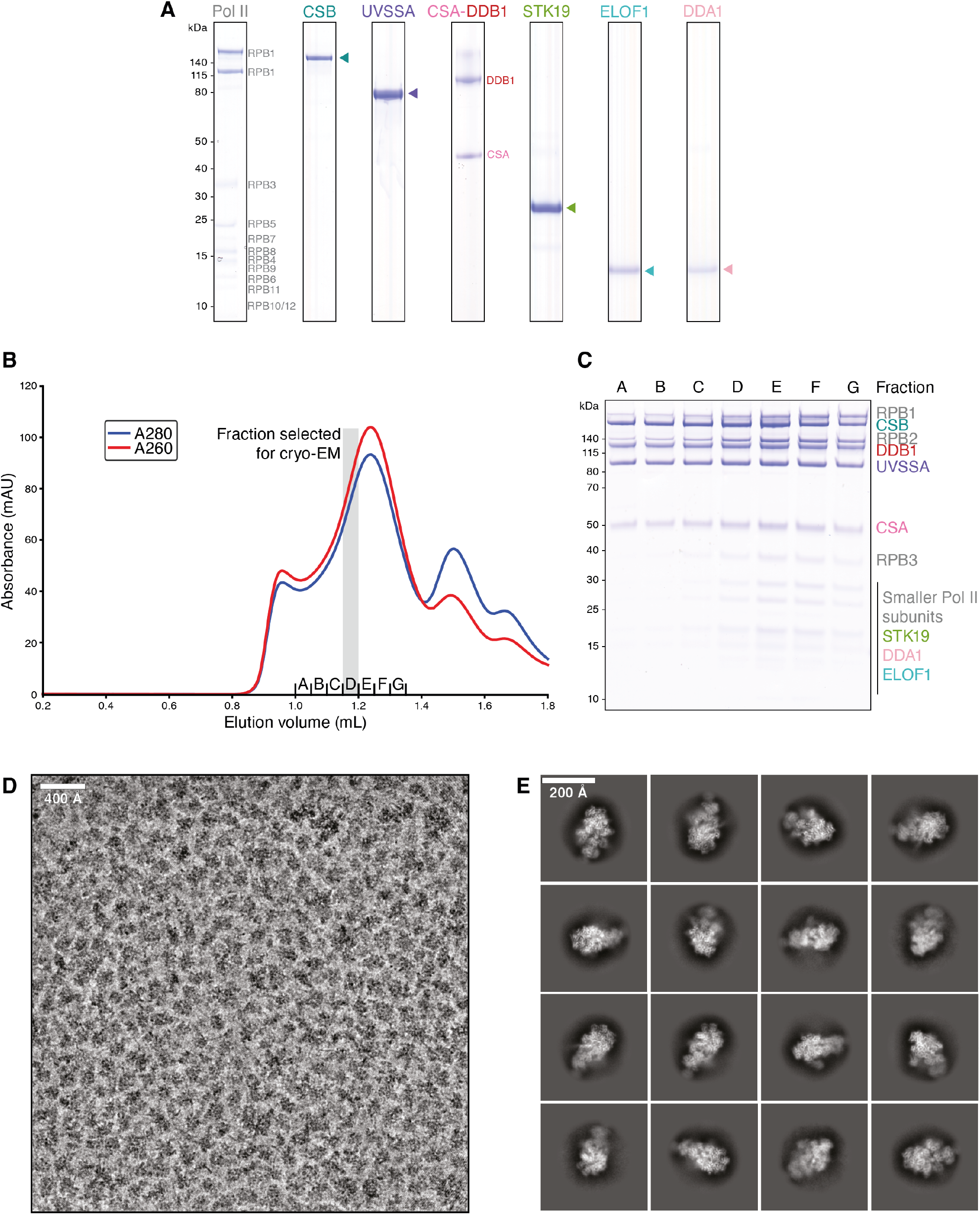
Complex formation and cryo-EM data analysis. **(A)** SDS-PAGE of purified *S. scrofa* RNA polymerase II and *H. sapiens* TC-NER factors, including STK19 and DDA1. **(B)** Chromatogram of TC-NER complex formation via size-exclusion chromatography. **(C)** SDS-PAGE of fractions from (B). **(D)** Representative micrograph from cryo-EM data collection. Scale bar, 400 Å. **(E)** 2D classes of TC-NER complex. Scale bar, 200 Å.

**Figure S5.**
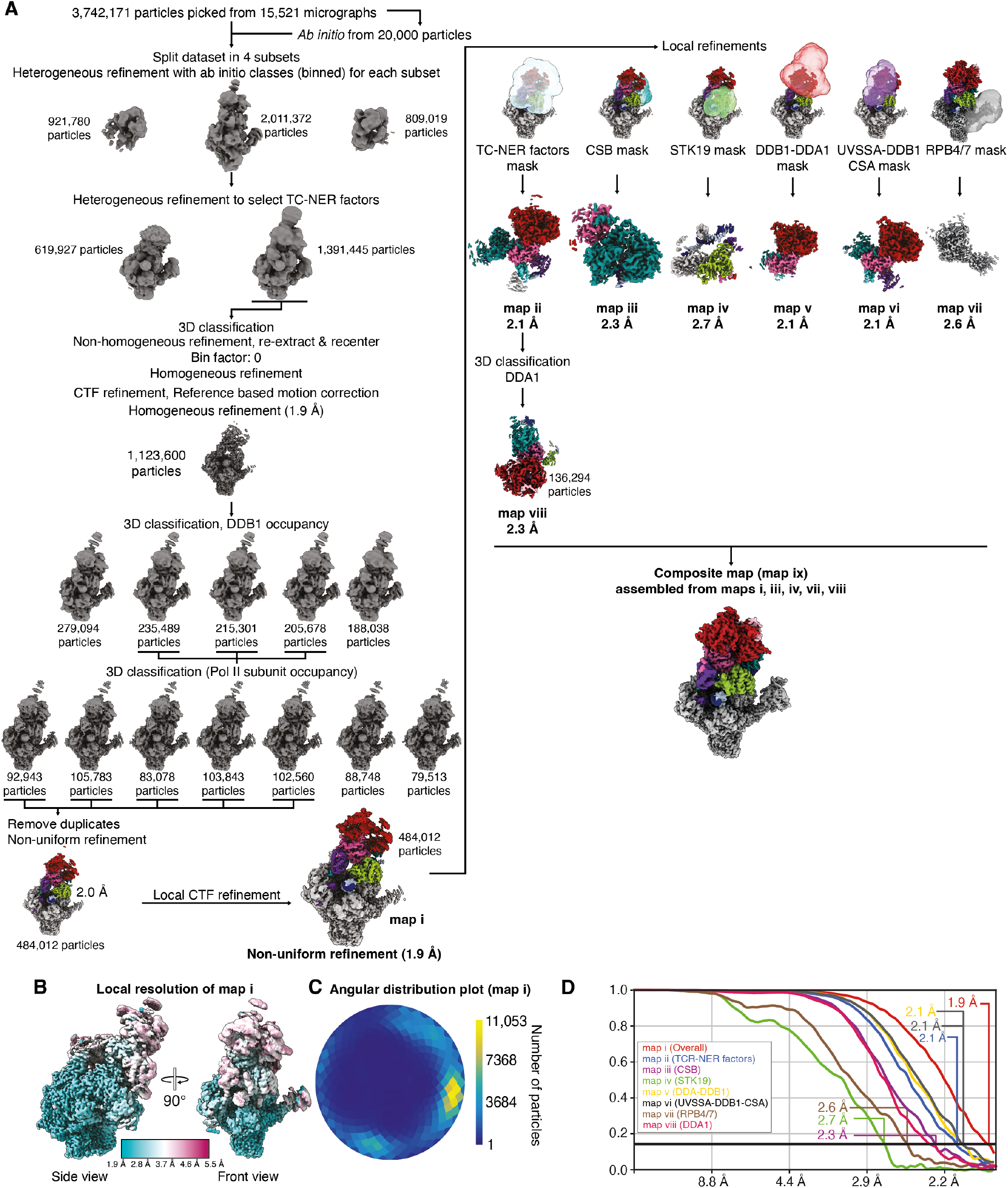
Classification tree and cryo-EM data analysis metrics. **(A)** Classification tree of cryo-EM data analysis. Particle numbers and resolutions are indicated. **(B)** Local resolution as colored on map i. **(C)** Angular distribution plot of particle assignment (map i). **(D)** Fourier shell correlation (FSC) curves of maps i-viii. FSC 0.143 criterion and achieved resolutions are indicated.

**Figure S6.**
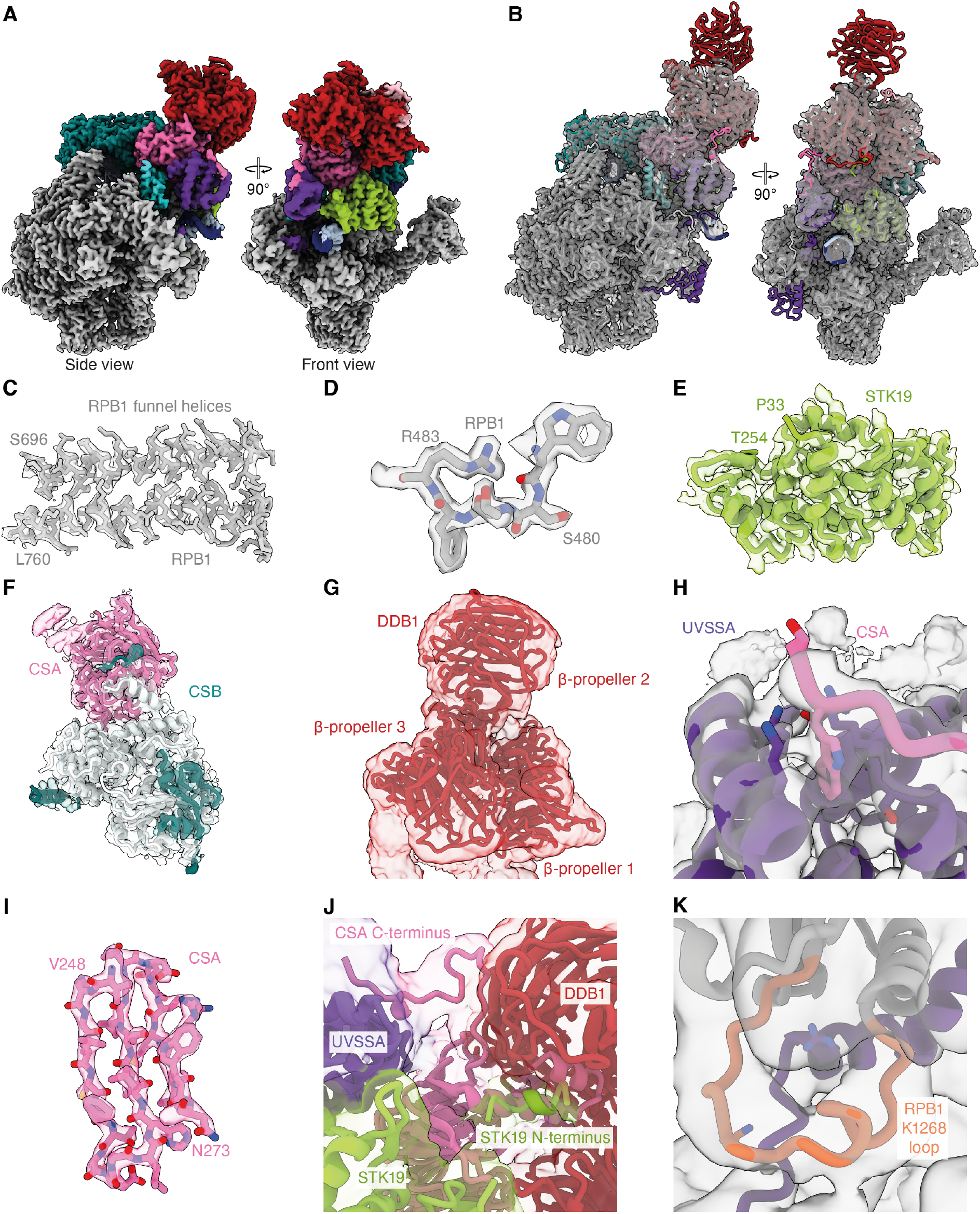
Cryo-EM data quality. **(A)** Coulomb potential map of RNA polymerase II TC-NER complex (composite map ix). **(B)** Coulomb potential map of RNA polymerase II TC-NER complex (grey, composite map ix) with fitted model. **(C)** RPB1 funnel helices with corresponding density (map i, sharpened). **(D)** Features of RPB1 resolves aromatic rings (map i, sharpened). **(E)** Coulomb potential map with corresponding model of STK19 (map iv, DeepEMhanced). **(F)** Coulomb potential map with corresponding model of CSA and CSB (map iii, DeepEMhanced). **(G)** Coulomb potential map with corresponding model of DDB1 (map ii, low-pass filtered). **(H)** Coulomb potential map with corresponding model of UVSSA and CSA C-terminus (map i, low-pass filtered). **(I)** Coulomb potential map with corresponding model of CSA residues 248-273 (map ii, sharpened. **(J)** Coulomb potential map with corresponding model of CSA C-terminus, STK19 N-terminus, UVSSA and DDB1 248-273 (map i, low-pass filtered). **(K)** Coulomb potential map with corresponding model of RPB1 K1268 loop and UVSSA C-terminus (map i, low-pass filtered).

**Figure S7.**
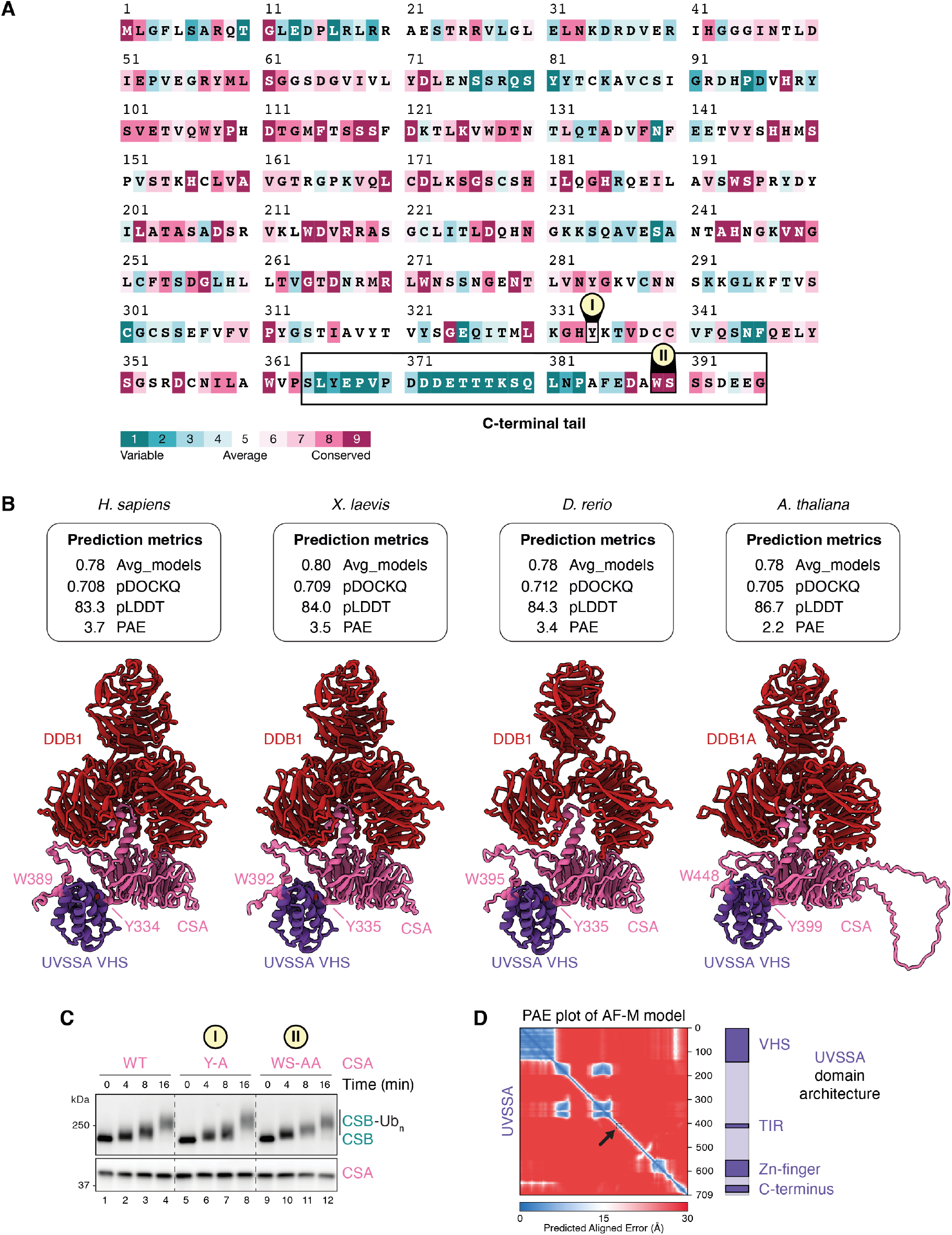
The interaction of the C-terminal tail of CSA with UVSSA is conserved. **(A)** Sequence conservation of human CSA calculated in ConSurf.^46^ CSA residues involved in the interaction with the UVSSA VHS domain as described in Figure 6A and 6B are highlighted. **(B)** AF-M structure prediction of full-length UVSSA in complex with CSA-DDB1 in the indicated organisms. For simplicity, only the VHS domain of UVSSA bound by CSA is depicted. Side chains of CSA residues interacting with UVSSA are shown. Confidence metrics quantify the interaction between UVSSA and the CSA-DDB1 heterodimer. **(C)** *In vitro* ubiquitination assay like the one shown in Figure 6C, except that UVSSA was replaced by human CSB and UBE2E1 was replaced by UBE2D2. Reaction products were blotted for CSA and CSB. CSB-Ub_n_, polyubiquitinated CSB. **(D)** PAE plot for the predicted full-length structure of human UVSSA and its domain architecture. The location of the TFIIH-interacting region (TIR) within the PAE plot is highlighted by an arrow.

**Table S1:** Confidence metrics for AlphaFold-Multimer predictions involving STK19. The attached table shows the confidence metrics SPOC (Structure Prediction and Omics Classifier), Avg_models, and pDOCKQ. See Methods.

**Table S2:**
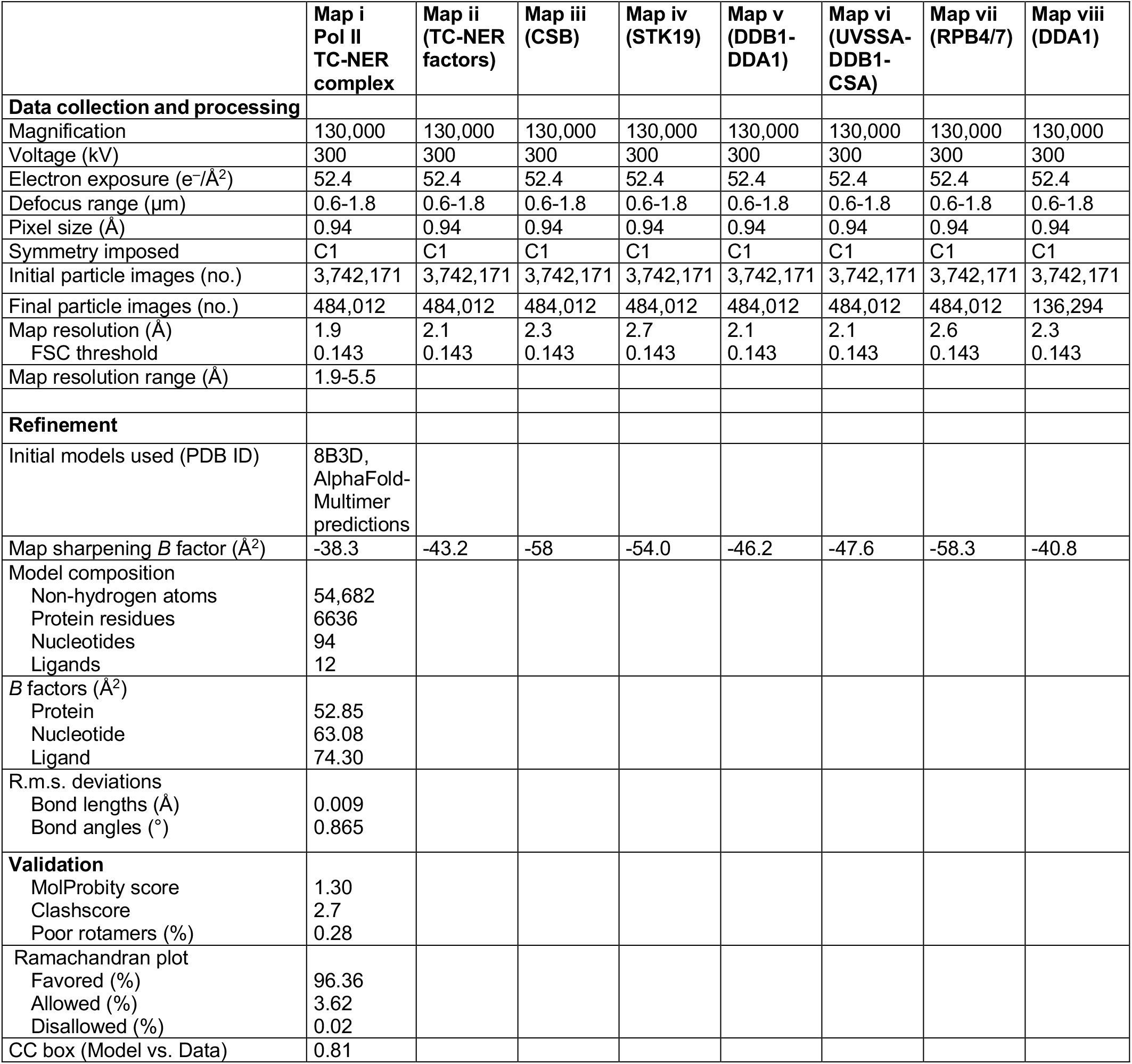
Cryo-EM data collection, refinement and validation statistics.

**Table S3:** Input structural models and model confidence.

| Subunit | Chain id(s) | Input model | Level of confidence |
| --- | --- | --- | --- |
| RPB1 | A | 8B3D | Atomic, RPB1 K1268 loop: secondary structure confidence |
| RPB2 | B | 8B3D | Atomic |
| RPB3 | C | 8B3D | Atomic |
| RPB4 | D | 8B3D | Secondary structure |
| RPB5 | E | 8B3D | Atomic |
| RPB6 | G | 8B3D | Atomic |
| RPB7 | F | 8B3D | Secondary structure |
| RPB8 | H | 8B3D | Atomic |
| RPB9 | I | 8B3D | Atomic |
| RPB10 | J | 8B3D | Atomic |
| RPB11 | K | 8B3D | Atomic |
| RPB12 | L | 8B3D | Atomic |
| ELOF1 | M | 8B3D | Atomic |
| Template DNA | T | 8B3D | Atomic |
| Non-template DNA | N | 8B3D | Atomic |
| RNA | P | 8B3D | Atomic |
| ERCC8/CSA | a | 8B3D | Atomic |
| ERCC6/CSB | b | AlphaFold2 | Atomic |
| UVSSA | c | 8B3D | Atomic, UVSSA Zn finger: clipped and rigid body docked, UVSSA C-terminus: clipped |
| DDB1 | d | AlphaFold-Multimer of DDB1/DDA1 model | Atomic, β-propeller 2: rigid body docked |
| DDA1 | e | AlphaFold-Multimer of DDB1/DDA1 model | Atomic |
| STK19 | f | AlphaFold-Multimer, *De novo* | Atomic, STK19 N-terminus: Backbone level confidence, no register confidence |

**Table S4:**
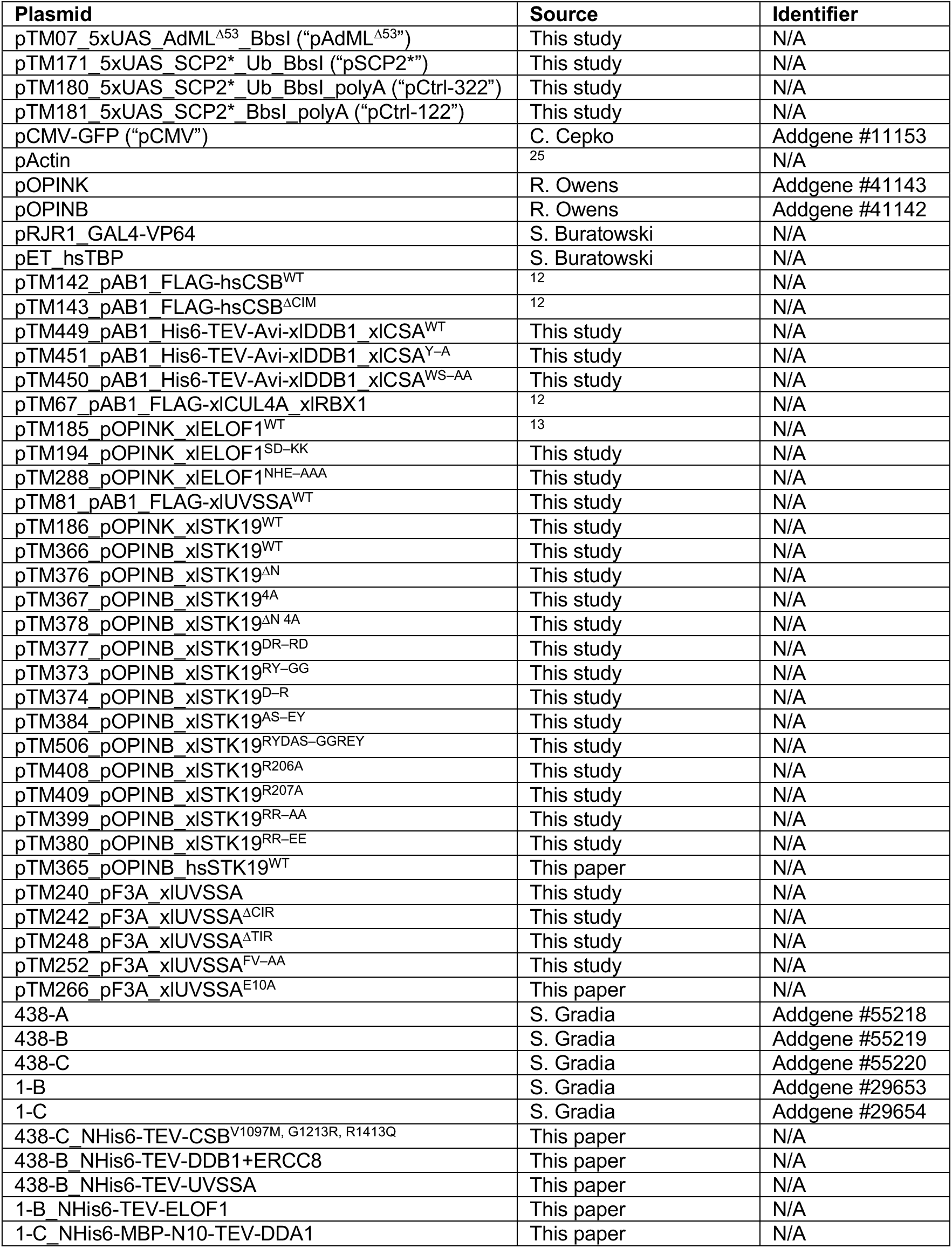
Plasmids.

**Table S5:**
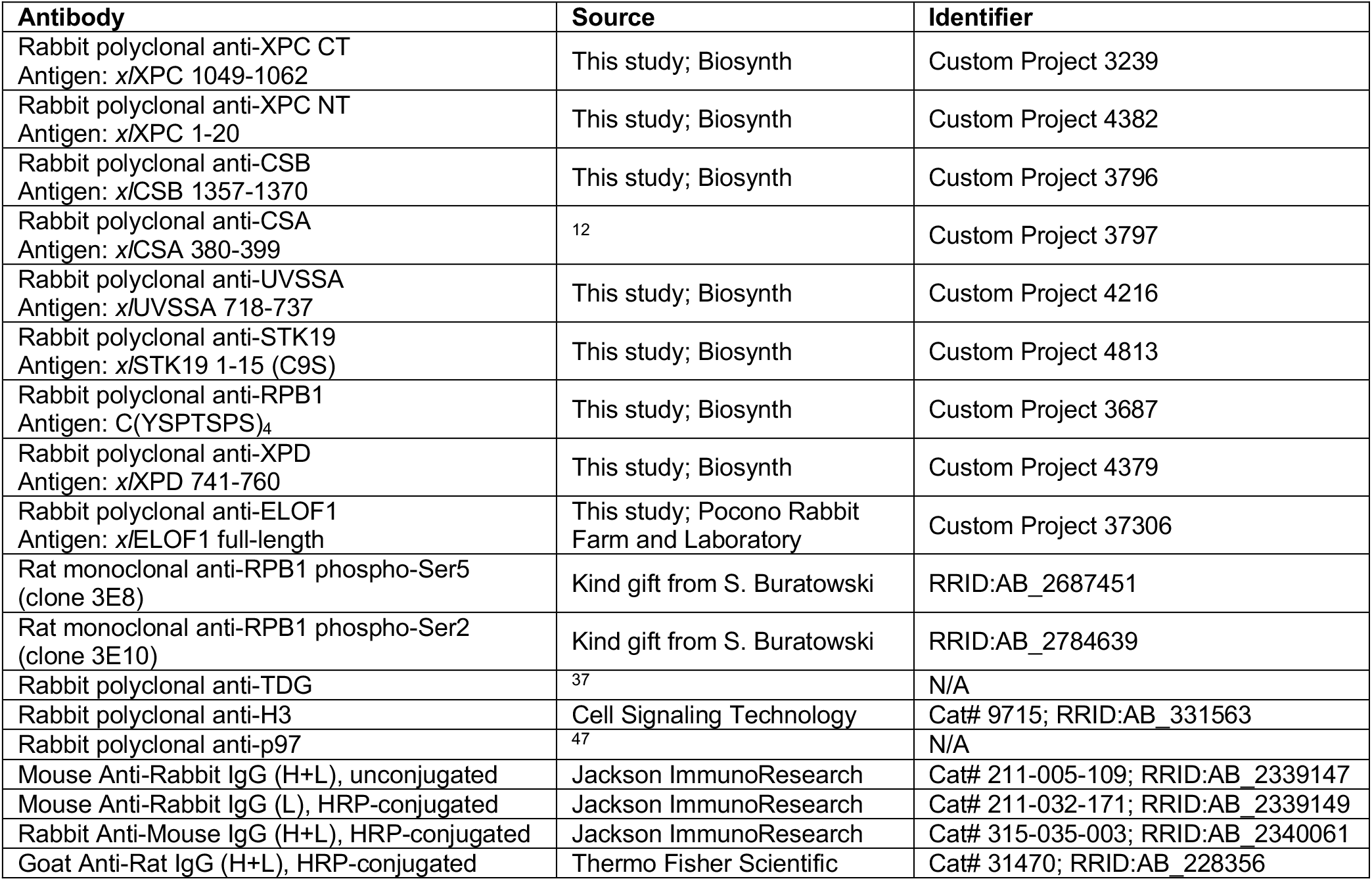
Antibodies.

